# A digital 3D reference atlas reveals cellular growth patterns shaping the Arabidopsis ovule

**DOI:** 10.1101/2020.09.19.303560

**Authors:** Athul Vijayan, Rachele Tofanelli, Soeren Strauss, Lorenzo Cerrone, Adrian Wolny, Joanna Strohmeier, Anna Kreshuk, Fred A. Hamprecht, Richard S. Smith, Kay Schneitz

## Abstract

A fundamental question in biology is how morphogenesis integrates the multitude of distinct processes that act at different scales, ranging from the molecular control of gene expression to cellular coordination in a tissue. Investigating morphogenesis of complex organs strongly benefits from three-dimensional representations of the organ under study. Here, we present a digital analysis of ovule development from *Arabidopsis thaliana* as a paradigm for a complex morphogenetic process. Using machine-learning-based image analysis we generated a three-dimensional atlas of ovule development with cellular resolution. It allows quantitative stage- and tissue-specific analysis of cellular patterns. Exploiting a fluorescent reporter enabled precise spatial determination of gene expression patterns, revealing subepidermal expression of *WUSCHEL*. Underlying the power of our approach, we found that primordium outgrowth progresses evenly, discovered a novel mode of forming a new cell layer, and detected a new function of *INNER NO OUTER* in restricting cell proliferation in the nucellus. Moreover, we identified two distinct subepidermal cell populations that make crucial contributions to ovule curvature. Our work demonstrates the expedience of a three-dimensional digital representation when studying the morphogenesis of an organ of complex cellular architecture and shape that eventually consists of 1,900 cells.

## Introduction

How organs attain their species-specific size and shape in a reproducible manner is an important question in biology. Tissue morphogenesis constitutes a multi-scale process that occurs in three dimensions (3D) plus time. Thus, quantitative cell and developmental biology must not only address molecular processes but also cellular and tissue-level properties. It necessitates the quantitative 3D analysis of cell size, cell shape, and cellular topology, an approach that has received much less attention (Boutros et al., 2015; Jackson et al., 2019).

Morphogenesis involves coordination of cellular behaviour between cells or complex populations of cells which may lead to emergent properties of the tissue that are not directly encoded in the genome (Coen et al., 2017; Coen and Rebocho, 2016; Gibson et al., 2011; Jackson et al., 2019). For example, plant cells are connected through their cell walls. The physical coupling of plant cells can cause mechanical stresses that may control tissue shape by influencing growth patterns and gene expression (Bassel et al., 2014; Hamant et al., 2008; Hervieux et al., 2016; Kierzkowski et al., 2012; Landrein et al., 2015; Louveaux et al., 2016; Sampathkumar et al., 2014; Sapala et al., 2018; Sassi et al., 2014; Uyttewaal et al., 2012). Concepts involving the minimization of mechanical stresses caused by differential growth within a tissue have been employed to explain morphogenesis of different plant organs with curved shapes (Lee et al., 2019; Liang and Mahadevan, 2011; Rebocho et al., 2017).

Developmental changes in the appearance of organs, such as leaves and sepals, are often assessed by focusing on the organ surface (Hong et al., 2016; Hervieux et al., 2016; Kierzkowski et al., 2019). This strategy, however, loses sight of internal cellular growth patterns. Cellular patterns in deeper tissue layers have classically been studied using 2D techniques with modern variations, for example in the study of the hypocotyl, relying on automated quantitative histology (Sankar et al., 2014). However, 2D analysis of cellular patterns can also result in misconceptions as was noticed for the early Arabidopsis embryo (Yoshida et al., 2014).

Such considerations make it obvious that quantitative and 3D cellular descriptions of the entire tissue under study are essential for a full understanding of tissue morphogenesis (Hong et al., 2018; Kierzkowski and Routier-Kierzkowska, 2019; Sapala et al., 2019). Digital 3D organs with complete cellular resolution represent a natural strategy to approach this endeavour. However, it remains an important challenge to generate such faithful digital representations. Substantial efforts in animal developmental biology did not achieve single-cell resolution (Lein et al., 2007, 2007; Rein et al., 2002; Lein et al., 2007; Dreyer et al., 2010; Asadulina et al., 2012; Anderson et al., 2019), except for *C. elegans* (Long et al., 2009) and the early embryo of the ascidian *Phallusia mammillata* (Guignard et al., 2020; Sladitschek et al., 2020). Model plants, such as *Arabidopsis thaliana*, are uniquely suited for this task. Plants feature a relatively small number of different cell types. Moreover, plant cells are large and immobile. As a consequence, one can often observe characteristic cell division patterns associated with the formation and organization of tissues and organs. So far, 3D digital organs were obtained for tissues of either small size or simple architecture and were incompatible with fluorescent stains (Bassel et al., 2014; Montenegro-Johnson et al., 2015; Pasternak et al., 2017; Schmidt et al., 2014; Yoshida et al., 2014).

The ovule is the major female reproductive organ of higher plants. It harbors the egg cell which is protected by two integuments, lateral determinate tissues that develop into the seed coat following fertilization. The Arabidopsis ovule has been established as an instructive model to study numerous aspects of tissue morphogenesis including primordium formation, the establishment of the female germ line, and integument formation (Chaudhary et al., 2018; Gasser and Skinner, 2019; Nakajima, 2018; Schmidt et al., 2015). The ovule exhibits an elaborate tissue architecture exemplified by its extreme curvature caused in part by the asymmetric growth of the two integuments. This property makes it ideal for addressing the complexity of morphogenetic processes. Qualitative descriptions of ovule development exist (Christensen et al., 1997; Robinson-Beers et al., 1992; Schneitz et al., 1995) but a quantitative cellular characterization is only available for the tissue that forms the germ line (Lora et al., 2017). To make the next step in the study of ovule morphogenesis therefore requires 3D digital ovules with cellular resolution and of all developmental stages.

Here, we constructed a canonical 4D digital atlas of Arabidopsis ovule development with cellular resolution. The atlas covers all stages from early primordium outgrowth to the mature pre-fertilization ovule, provides quantitative information about various cellular parameters, and provides a proof-of-concept analysis regarding gene expression in 3D with cellular resolution. Our quantitative phenotypic analysis revealed a range of novel aspects of ovule morphogenesis and a new function for the regulatory gene *INNER NO OUTER*.

## Results

Arabidopsis ovules become apparent as finger-like protrusions that emanate from the placental tissue of the carpel (Robinson-Beers et al., 1992; Schneitz et al., 1995) (Fig. 1A). The ovule is a composite of three clonally distinct radial layers (Jenik and Irish, 2000; Schneitz et al., 1995). Thus, its organization into L1 (epidermis), L2 (first subepidermal layer) and L3 (innermost layer) follows a general principle of plant organ architecture (Satina et al., 1940). Following primordium formation, three proximal-distal (PD) pattern elements can be recognized: the distal nucellus, central chalaza and proximal funiculus, respectively Fig. 1A,B). The nucellus produces the megaspore mother cell (MMC), a large L2-derived cell that eventually undergoes meiosis. Only one of the meiotic products, the functional megaspore, survives and continues development. It develops into the eight-nuclear, seven-celled haploid embryo sac. The embryo sac, or female gametophyte, carries the actual egg cell. The chalaza is characterized by two integuments that initiate at its flanks. The two sheet-like integuments are determinate lateral organs of epidermal origin that undergo planar or laminar growth (Jenik and Irish, 2000; Schneitz et al., 1995; Truernit and Haseloff, 2008). The integuments grow around the nucellus in an asymmetric fashion eventually forming a hood-like structure and contributing to the curved shape (anatropy) of the mature ovule. Each of the two integuments initially forms a bi-layered structure of regularly arranged cells. Eventually, the inner integument forms a third layer. The outer integument consists of two cell layers throughout its development. The two integuments leave open a small cleft, the micropyle, through which a pollen tube can reach the interior of the ovule (Fig. 1A,C). The funiculus represents a stalk-like structure that carries the vasculature and connects the ovule to the placenta. Ovules eventually orient along the long axis of the gynoecium with the micropyle facing towards the stigma (gynapical side) while the opposite side of the ovule faces away from the stigma (gynbasal side) (Fig. 1B).

**Figure 1.**
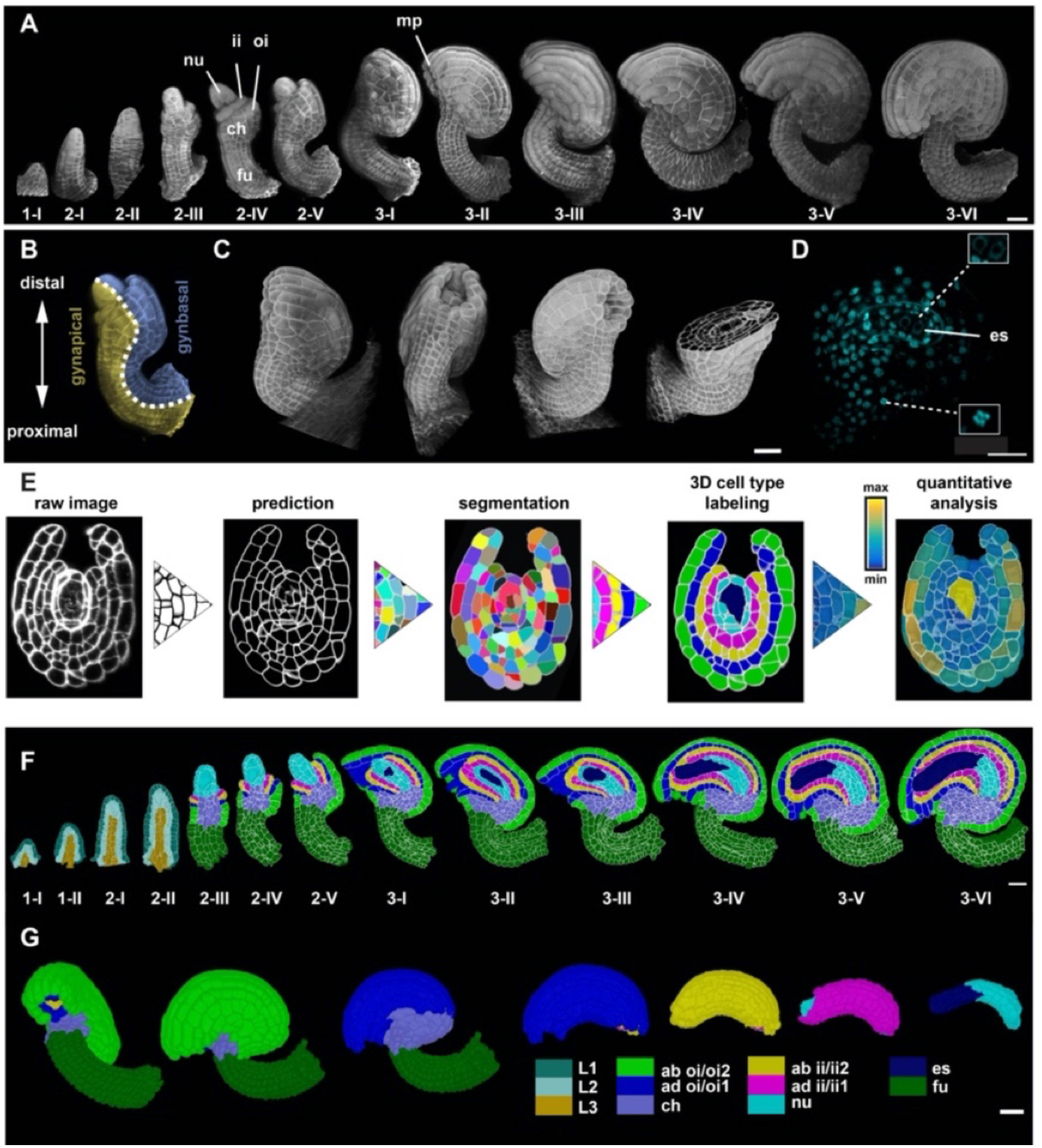
Stage-specific 3D digital ovules with cellular and tissue resolution. (A) 3D rendering of confocal z-stacks of SR2200 stained cell walls of ovule depicting ovule development from initiation at stage 1-I to maturity 3-VI. (B) The different polarities of the ovule: the proximal-distal axis and the gynapical-gynbasal axis are indicated. (C) 3D rendering of confocal z-stacks with multi-view of an ovule depicting the quality of raw microscopic image. (D) Mid-section clip plane from the TO-PRO-3 channel displaying a two-nuclear embryo sac and mitotic nuclei. (E) Pipeline generating 3D digital ovules: raw data, PlantSeg cell contour prediction, 3D GASP segmentation, cell type annotation and quantitative analysis. (F) Mid-sagittal section of ovules from stage 1-I to 3-VI showing the cell type organization in wild-type ovules. Stages 1-I to 2-II includes radial L1, L2, L3 labelling. From stage 2-III individual cell type labels are assigned according to the specific tissue. (G) 3D view of a mature ovule with cell type labels. The inner tissues are extracted from the 3D ovule after removing the overlying tissues and visualized separately. Different colors represent different tissue type labels. ii1/ii2, oi1/oi2 designate the integument layers as described in (Beeckman et al., 2000). Abbreviations: ab, abaxial; ad, adaxial; ch, chalaza; es, embryo sac; fu, funiculus; ii, inner integument; mp, micropyle; nu, nucellus; oi, outer integument. Scale bars: 20 μm.

### Generating stage-specific 3D digital ovules with cellular resolution

Ovules are buried within the gynoecium and faithful live imaging of Arabidopsis ovule development is not feasible, except for a short period of time and with a focus on a given cell (Tofanelli et al., 2019; Valuchova et al., 2020). Thus, we resorted to imaging cohorts of fixed specimens. We obtained z-stacks of optical sections of fixed and cleared ovules at different stages by laser scanning confocal microscopy (CLSM). The image stacks were further handled using MorphoGraphX (MGX) software (Barbier de Reuille et al., 2015; Strauss et al., 2019). The imaging method has recently been described in detail (Tofanelli et al., 2019). In short, we dissected and fixed ovules of different stages, cleared the ovules using ClearSee (Kurihara et al., 2015) and simultaneously stained the cell wall and nuclei using the fluorescent stains SR2200 (Musielak et al., 2015) and TO-PRO-3 iodide (TO-PRO-3) (Bink et al., 2001; Van Hooijdonk et al., 1994), respectively (Fig. 1C,D).

We processed the raw 3D datasets using PlantSeg, a deep learning pipeline for 3D segmentation of dense plant tissue at cellular resolution (Wolny et al., 2020) (Fig. 1E). The pipeline includes two major steps: cell wall stain-based cell boundary prediction performed by a convolutional neural network (CNN) and 3D cell segmentation based on the respective cell boundary predictions. Even after extensive optimization (see Materials and Methods), the procedure still resulted in some mistakes. We found that two distinct groups of cells represented the main sources of error. The first group included the MMC and its direct lateral neighbors at stages 2-III to 2-V. The second group encompassed the cells of the late embryo sac (stages 3-V/ 3-VI) (Tofanelli et al., 2019). We believe the reason for these errors lies in poor staining with SR2200 and could be due to their cell walls being particularly thin or of a biochemical composition interfering with staining. Poor staining of the embryo sac cells could be explained by the observation that they exhibit only partially formed cell walls (Mansfield et al., 1991). An additional minor source of errors related to remaining general segmentation mistakes which were partly manually corrected. If the remaining segmentation mistakes exceeded a given threshold, the ovule was excluded from further analysis (see below).

Using this methodology we generated a high-quality 3D image dataset consisting of 158 wild-type ovules of the accession Columbia (Col-0, ≥ 10 samples per stage). We selected ovules devoid of apparent segmentation errors for stages 1 to 2-II and 3-I to 3-IV. For stages 2-III to 2-V we included ovules containing no more than five under-segmented cells in the region occupied by the MMC and its lateral neighboring cells (≤ 10% of nucellar cells). Regarding mature ovules (stages 3-V/3-VI) we included ovules devoid of apparent segmentation errors in the sporophytic tissue.

### Cell-type labelling of 3D digital ovules with cellular resolution

Following the generation of 3D cell meshes we added specific labels to individual cells, thereby describing tissue type, such as radial cell layers (L1, L2, L3), nucellus, internal tissue of the chalaza, inner or outer integument, or the funiculus (see Table S1 for cell types). Staining with TO-PRO-3 also allowed the identification of cells undergoing mitosis (Fig. 1D). Available computational pipelines for near-automatic, geometry-based cell type identification (Montenegro-Johnson et al., 2015; Montenegro-Johnson et al., 2019; Schmidt et al., 2014) failed to provide reasonably good and consistent results. This was likely due to the ovule exhibiting a more complex tissue architecture. We therefore performed cell type labelling by a combination of semi-automated and manual cell type labelling. The combined efforts resulted in a reference set of 158 hand-curated 3D digital ovules of the wild-type Col-0 accession. They feature cellular resolution, cover all stages, and include annotated cell types and cellular features (Fig. 1F).

### Overall assessment of ovule development

The different stages of Arabidopsis ovule development were defined previously (Schneitz et al., 1995). Throughout this work we used a more precise definition of the subdivision of stage 1 into stage 1-I and 1-II. The subdivision was based on the first appearance of the signal of a reporter for *WUSCHEL* expression which became robustly apparent when ovule primordia consisted of 50 cells (see below). We first determined the average number of cells per ovule and stage (Fig. 2A,B). We counted the cells per ovule at different stages and found an incremental increase in cell number for every consecutive stage of ovule development until ovules at stage 3-VI exhibited an average of about 1900 cells (1897 ± 179.9 (mean ± SD)) (Table 1). We also assessed the mean volume per ovule for each stage by summing up the cell volumes of all cells in a given ovule (Fig. 2A,C) (Table 1). We measured a mean total volume for ovules at stage 3-VI of about 5×10^5^ μm^3^ (4.9×10^5^ ± 0.7 ×10^5^).

**Figure 2.**
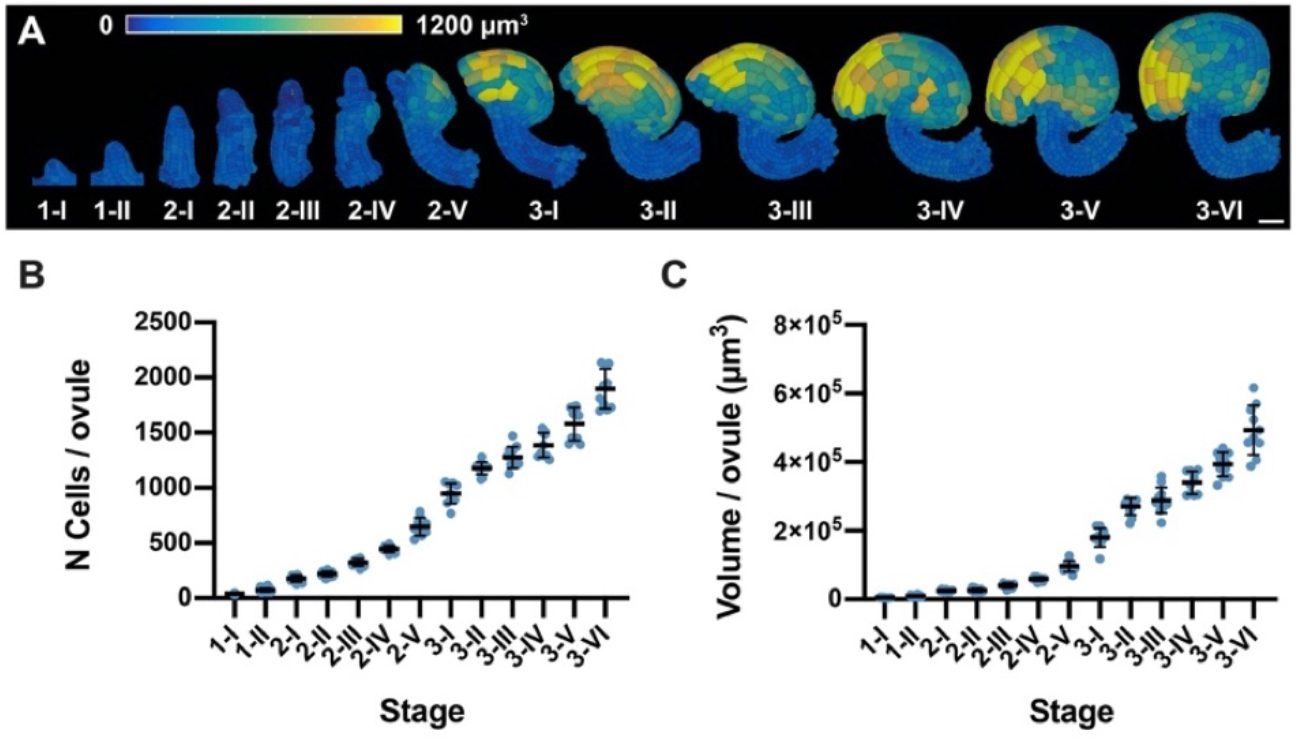
Ovule developmental stages and overall growth patterns. (A) 3D cell mesh view of wild-type ovules at different stages displaying heatmaps of cell volume ranging from 0 to 1200 μm^3^. (B, C) Plots depicting the total number of cells and total volume of individual ovules from early to late stages of development, respectively. Mean ± SD is shown. Scale bar: 20 μm.

**Table 1.**
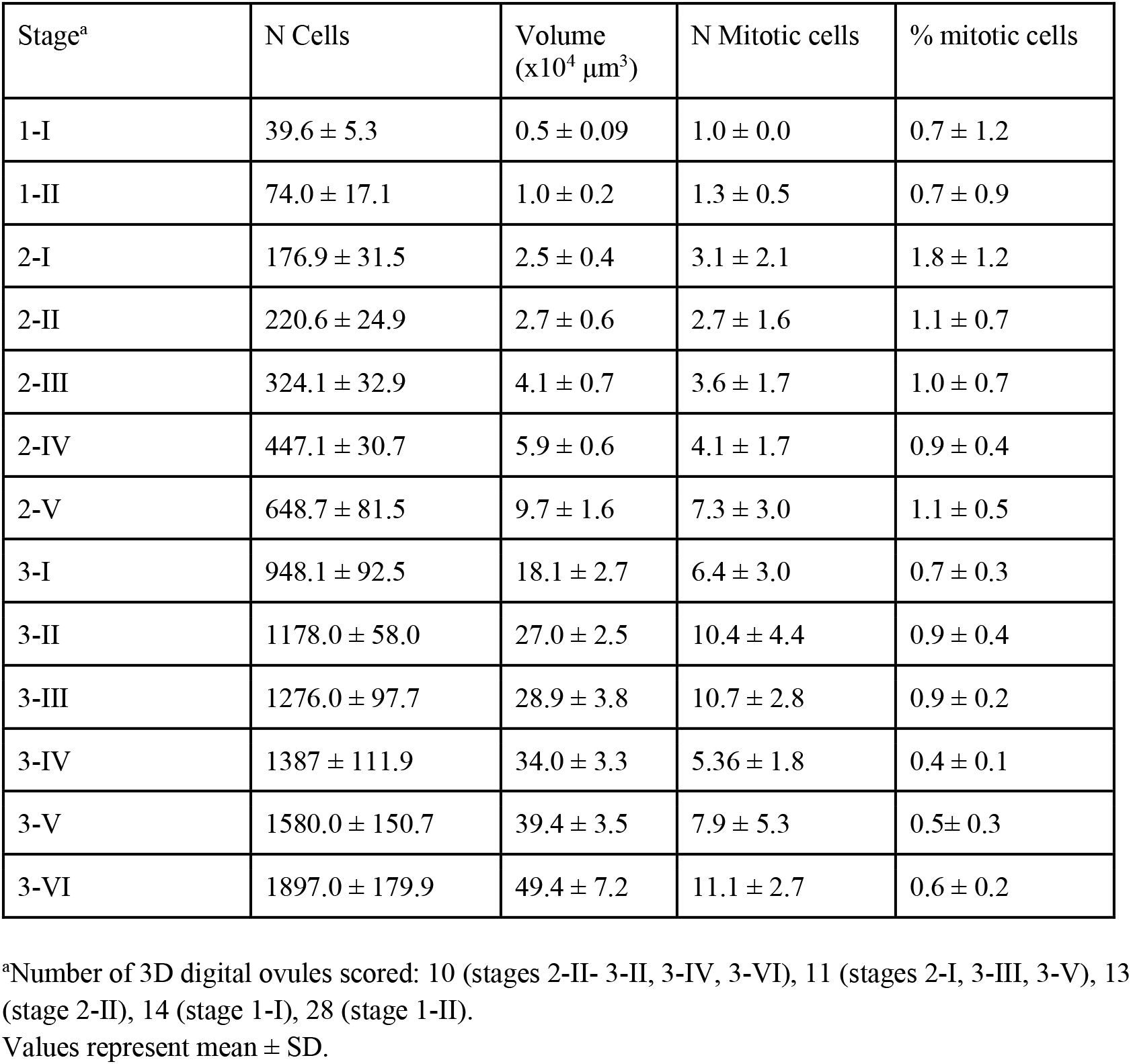
Cell numbers and total volumes of ovules at different stages.

| Stage<sup>a</sup> | N Cells | Volume<br>($\times 10^4 \mu\text{m}^3$) | N Mitotic cells | % mitotic cells |
| --- | --- | --- | --- | --- |
| 1-I | 39.6 ± 5.3 | 0.5 ± 0.09 | 1.0 ± 0.0 | 0.7 ± 1.2 |
| 1-II | 74.0 ± 17.1 | 1.0 ± 0.2 | 1.3 ± 0.5 | 0.7 ± 0.9 |
| 2-I | 176.9 ± 31.5 | 2.5 ± 0.4 | 3.1 ± 2.1 | 1.8 ± 1.2 |
| 2-II | 220.6 ± 24.9 | 2.7 ± 0.6 | 2.7 ± 1.6 | 1.1 ± 0.7 |
| 2-III | 324.1 ± 32.9 | 4.1 ± 0.7 | 3.6 ± 1.7 | 1.0 ± 0.7 |
| 2-IV | 447.1 ± 30.7 | 5.9 ± 0.6 | 4.1 ± 1.7 | 0.9 ± 0.4 |
| 2-V | 648.7 ± 81.5 | 9.7 ± 1.6 | 7.3 ± 3.0 | 1.1 ± 0.5 |
| 3-I | 948.1 ± 92.5 | 18.1 ± 2.7 | 6.4 ± 3.0 | 0.7 ± 0.3 |
| 3-II | 1178.0 ± 58.0 | 27.0 ± 2.5 | 10.4 ± 4.4 | 0.9 ± 0.4 |
| 3-III | 1276.0 ± 97.7 | 28.9 ± 3.8 | 10.7 ± 2.8 | 0.9 ± 0.2 |
| 3-IV | 1387 ± 111.9 | 34.0 ± 3.3 | 5.36 ± 1.8 | 0.4 ± 0.1 |
| 3-V | 1580.0 ± 150.7 | 39.4 ± 3.5 | 7.9 ± 5.3 | 0.5 ± 0.3 |
| 3-VI | 1897.0 ± 179.9 | 49.4 ± 7.2 | 11.1 ± 2.7 | 0.6 ± 0.2 |

<sup>a</sup>Number of 3D digital ovules scored: 10 (stages 2-II- 3-II, 3-IV, 3-VI), 11 (stages 2-I, 3-III, 3-V), 13 (stage 2-II), 14 (stage 1-I), 28 (stage 1-II).
Values represent mean ± SD.

### Tissue-specific growth patterns along the PD axis following primordium formation

Next we addressed the question if there exist tissue-specific growth differences. To this end we counted cell numbers and total volumes in the nucellus, embryo sac, chalaza, funiculus and the epidermal cells of the two integuments of stage 2-III to 3-VI digital ovules (Figures 3 and 4) (Table 2). We observed that cell numbers in the nucellus stayed roughly constant from stage 2-III up to 3-III (Fig. 3A). At stage 3-I the average cell number per nucellus was 66.9 ± 13.4. The results suggest that little if any cell proliferation takes place during these stages in the nucellus. Starting with stage 3-IV, however, cell numbers increased and we found an average cell number of 85.9 ± 17.2 at stage 3-VI hinting at somewhat elevated cell proliferation during these latter stages. This pattern was mirrored by a 2.2-fold increase in the average volume of the nucellus across different stages (excluding the developing embryo sac) (Fig. 3B) (stage 2-III: 0.6 × 10^4^ μm^3^ ± 0.1 10^4^ μm^3^; stage 3-VI: 1.3 × 10^4^ μm^3^ ± 0.2 × 10^4^ μm^3^). Beginning from stage 3-IV we failed to detect nucellar cells at the very micropylar (distal) end of the developing embryo sac (Fig. 1F). This observation confirmed previous observations (Schneitz et al., 1995) and raised the question what happened to these distal nucellar cells. Although we cannot exclude that a small number of cells becomes crushed we did not observe such events in this region. Thus, as the average cell number per nucellus stays constant or even increases slightly we propose that the growing embryo sac “pushes away” some distal nucellar cells.

**Figure 3.**
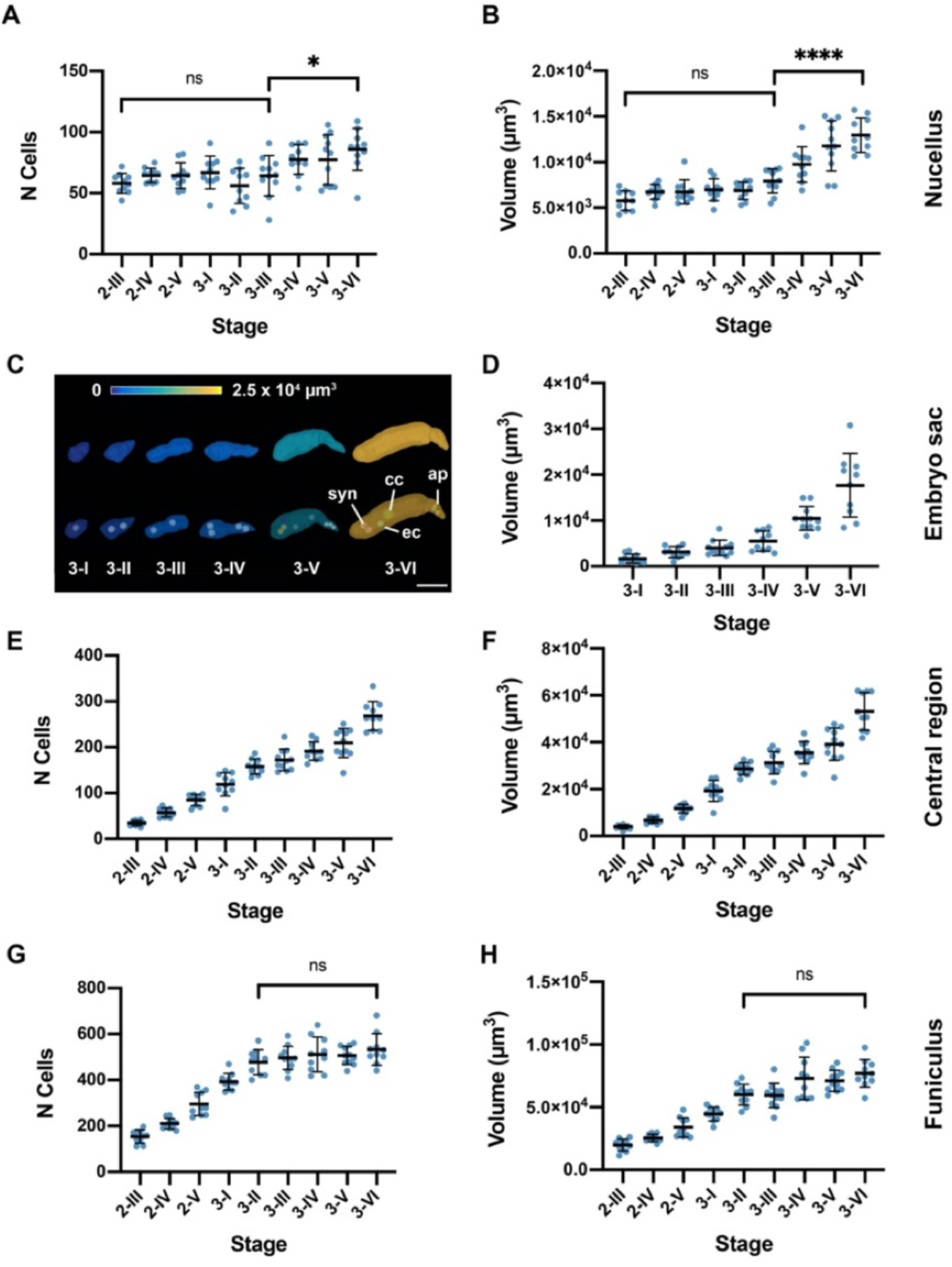
Tissue-specific quantitative analysis. (A, B) Plots depicting the number of cells and volume at different developmental stages of the nucellus, respectively. (C) 3D mesh of the embryo sac, from stage 3-I to stage 3-IV, extracted from 3D ovule cell meshes. The volume is represented as a heat map. (D) Plot depicting the embryo sac volume from individual ovule datasets at different stages. (E, F) Plots showing the number of cells and volume of the central region, respectively. (G, H) Plots depicting the number of cells and volume of the funiculus, respectively. Data points indicate individual ovules. Mean ± SD are represented as bars. Asterisks represent statistical significance (ns, P ≥ 0.5; *, P < 0.05; ****, P<0.0001; Student’s t test). Scale bar: 20 μm.

**Figure 4.**
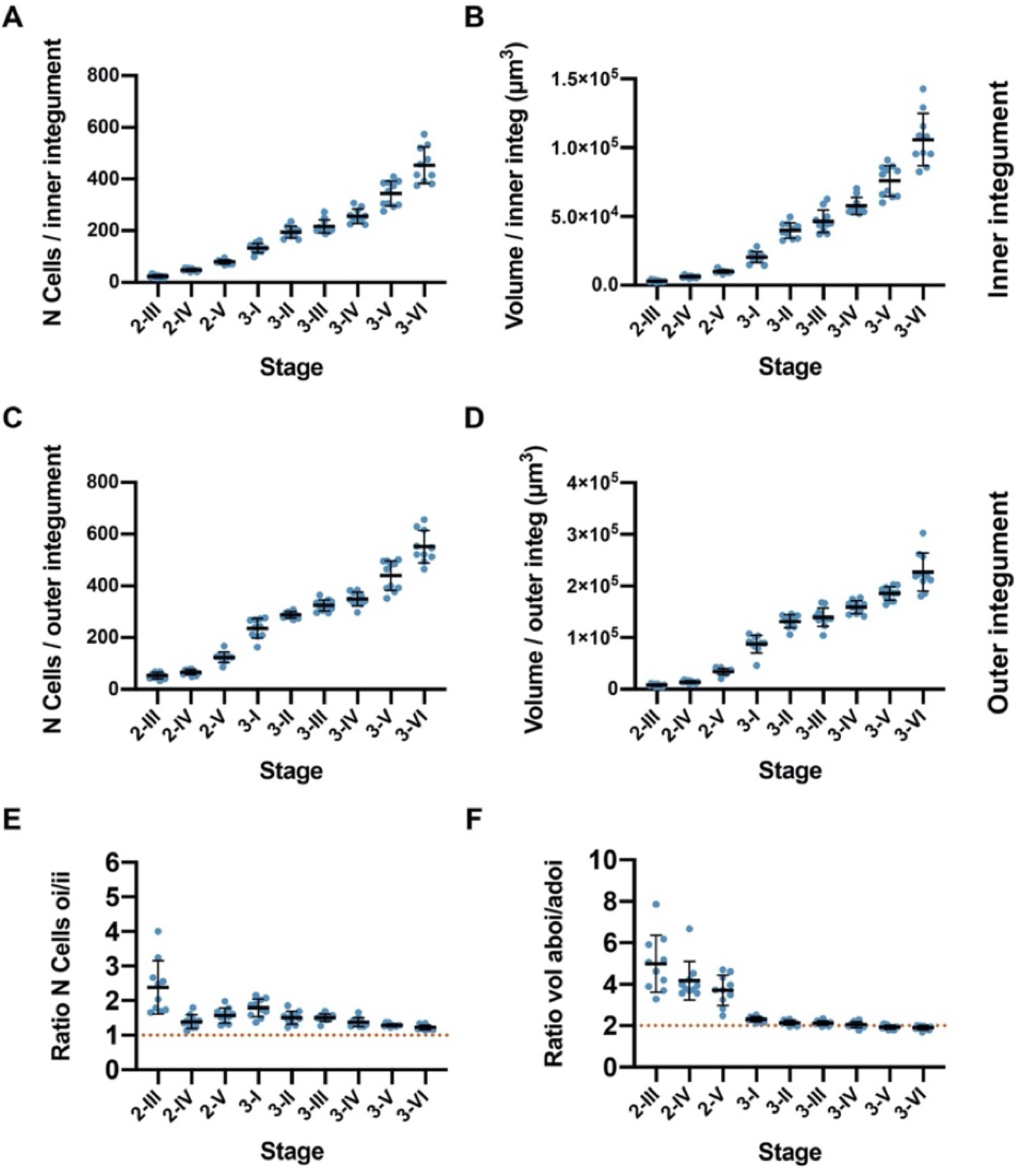
Quantitative analysis of cellular patterns in the integuments. (A, B) Plots indicating the number of cells and volume of the inner integument from stage 2-III to stage 3-VI. (C, D) Plots indicating the number of cells and volume of the outer integument from stage 2-III to 3-VI stages. (E, F) Plot showing the ratio between the number of cells and tissue volume of the outer and inner integument. Data points indicate individual ovules. Mean ± SD are represented as bars.

**Table 2.** Cell numbers and total volumes of the major ovule tissues.

|  | Tissue |  |  |  |  |  |  |  |  |  |
| --- | --- | --- | --- | --- | --- | --- | --- | --- | --- | --- |
| Stage<sup>a</sup> | Nucellus |  | Central Region |  | Inner Integument |  | Outer Integument |  | Funiculus |  |
| | N cells | Volume<br>($\times 10^4 \mu\text{m}^3$) | N cells | Volume<br>($\times 10^4 \mu\text{m}^3$) | N cells | Volume<br>($\times 10^4 \mu\text{m}^3$) | N cells | Volume<br>($\times 10^4 \mu\text{m}^3$) | N cells | Volume<br>($\times 10^4 \mu\text{m}^3$) |
| 2-III | 58.1 ± 8.1 | 0.6 ± 0.1 | 35.0 ± 5.7 | 0.4 ± 0.08 | 23.5 ± 5.5 | 0.30 ± 0.07 | 53.8 ± 12.1 | 0.8 ± 0.2 | 153.7 ± 28.5 | 2.0 ± 0.5 |
| 2-IV | 64.6 ± 5.9 | 0.7 ± 0.1 | 57.6 ± 9.7 | 0.7 ± 0.01 | 48.1 ± 5.9 | 0.62 ± 0.08 | 66.6 ± 10.6 | 1.4 ± 0.3 | 210.2 ± 23.6 | 2.5 ± 0.3 |
| 2-V | 64.5 ± 10.5 | 0.7 ± 0.1 | 85.0 ± 12.2 | 1.2 ± 0.2 | 79.3 ± 8.4 | 1.0 ± 0.1 | 124.0 ± 20.2 | 3.4 ± 0.6 | 295.9 ± 50.0 | 3.4 ± 0.8 |
| 3-I | 66.9 ± 13.4 | 0.7 ± 0.1 | 118.9 ± 25.4 | 1.9 ± 0.4 | 132.6 ± 18.2 | 2.04 ± 0.4 | 235.9 ± 37.6 | 8.8 ± 1.7 | 392.4 ± 36.4 | 4.5 ± 0.6 |
| 3-II | 56.0 ± 14.4 | 0.7 ± 0.1 | 158.1 ± 16.6 | 2.9 ± 0.3 | 194.7 ± 22.9 | 4.0 ± 0.6 | 289.1 ± 12.6 | 13.2 ± 1.3 | 477.1 ± 54.2 | 6.0 ± 0.8 |
| 3-III | 64.4 ± 16.6 | 0.8 ± 0.1 | 172.2 ± 23.4 | 3.1 ± 0.5 | 216.8 ± 25.4 | 4.7 ± 0.8 | 324.5 ± 21.4 | 13.9 ± 1.8 | 496.2 ± 50.6 | 6.0 ± 1.0 |
| 3-IV | 77.7 ± 12.2 | 1.0 ± 0.2 | 191.3 ± 20.1 | 3.6 ± 0.5 | 255.8 ± 28.4 | 5.8 ± 0.6 | 349.6 ± 26.6 | 15.9 ± 1.3 | 511.1 ± 75.8 | 7.3 ± 1.7 |
| 3-V | 77.4 ± 20.4 | 1.2 ± 0.3 | 209.2 ± 31.7 | 3.9 ± 0.7 | 343.5 ± 48.3 | 7.6 ± 1.1 | 439.8 ± 57.0 | 18.6 ± 1.3 | 506.8 ± 40.1 | 7.1 ± 0.8 |
| 3-VI | 85.9 ± 17.2 | 1.3 ± 0.2 | 268.5 ± 31.4 | 5.3 ± 0.8 | 453.9 ± 69.8 | 10.6 ± 1.9 | 551.6 ± 62.7 | 22.7 ± 3.7 | 533.0 ± 68.6 | 7.7 ± 1.1 |

<sup>a</sup>Number of 3D digital ovules scored: 10 (stages 2-II- 3-II, 3-IV, 3-VI), 11 (stages 3-III, 3-V).1345 Values represent mean ± SD

Following the formation of the functional megaspore coenocytic embryo sac development starts at stage 3-I. Three rounds of mitoses followed by cellularization eventually result in the typical eight-nuclear, seven-celled *Polygonum*-type of embryo sac found at stage 3-VI. The mono-nuclear embryo sac features a volume of 0.2 × 10^4^ μm^3^ ± 0.1 × 10^4^ μm^3^. At stage 3-VI we observed a total volume of 1.8 × 10^4^ μm^3^ ± 0.7 × 10^4^ μm^3^, a nine-fold increase (Fig 3C).

The central region of the ovule, classically known as chalaza, is of complex composition (see below). Here, we discriminate between the epidermis-derived cells of the two integuments and the internal cells of the central region encapsulated by the epidermis and first focus on the latter. We found a steady increase in the average internal cell number per central region from stage 2-III (35.0 ± 5.7) up to stage 3-VI (268.5 ± 31.4) (Fig. 3E,F) (Table 2). We also observed a steady increase in volume of the internal central region (stage 2-III: 0.4 × 10^4^ μm^3^ ± 0.1 10^4^ μm^3^; stage 3-VI: 5.3 × 10^4^ μm^3^ ± 0.8 × 10^4^ μm^3^). Thus, cell numbers increased 7.7 fold while tissue volume increased 13.3 fold over the scored stages.

Regarding cell number in the funiculus we observed a steady increase in the average cell number per funiculus from stage 2-III until the average cell number per funiculus reached 477.1 ± 54.2 at stage 3-II (Fig. 3G) (Table 2). This value stayed about constant throughout the later stages although we observed a minor and statistically insignificant increase to 533.0 ± 68.6 at stage 3-VI. We further observed an average volume at stage 3-VI of 7.7 × 10^4^ μm^3^ ± 1.1 × 10^4^ μm^3^ (Fig. 3H). The data suggest that there is very little if any growth in the funiculus after stage 3-II.

Next we assessed the number of epidermis-derived cells of the two integuments (Fig. 4) (Table 2). Both integuments showed continuous increases in cell number and integument volume for stages 2-III to 3-VI. We observed a 19.3-fold increase in cell number from stage 2-III to 3-VI. At stage 2-III the inner integument featured an average cell number of 23.5 ± 5.5 and at stage 3-VI of 453.9 ± 69.8 (Fig. 4A). Regarding the volume increase of the inner integument we observed an 35.3-fold increase in inner integument volume with an average volume of 0.3 × 10^4^ μm^3^ ± 0.07 × 10^4^ μm^3^ at stage 2-III and 10.6 × 10^4^ μm^3^ ± 1.9 × 10^4^ μm^3^ at stage 3-VI (Fig. 4B). For the outer integument we observed a 10.3-fold increase in cell number with an average number of cells of 53.8 ± 12.1 at stage 2-III and of 551.6 ± 62.7 at stage 3-VI (Fig. 4C). Regarding the volume increase of the outer integument we observed a 28.4-fold increase in volume with an average volume of 0.8 × 10^4^ μm^3^ ± 0.02 × 10^4^ μm^3^ at stage 2-III and 22.7 × 10^4^ μm^3^ ± 3.7 × 10^4^ μm^3^ at stage 3-VI (Fig. 4D). We also investigated the ratio of number of cells and volume of the outer integument versus the inner integument (Fig. 4E,F). The data revealed that the outer integument always carried more cells than the inner integument, although the difference was relatively small throughout the various stages with stage 3-VI showing a ratio of 1.2. By contrast, the volume of the outer integument was always more than twice the volume of the inner integument. At stage 3-I in particular we noticed a 4.3-fold larger volume of the outer integument. This value slowly decreased and at stage 3-VI we determined a ratio of 2.2.

### Differential growth patterns contribute to ovule development

To gain more insight into the growth dynamics of ovule morphogenesis we estimated stage-specific growth rates by taking the ratio between the mean total volume or mean total cell number of a stage and the corresponding values of the preceding stage (Fig. 5A,B). We observed that the volume increase was highest during primordium formation followed by a noticeable drop during stage 2-I. Growth increased again during stages 2-II to 3-I followed by another drop and a comparably flat curve during the rest of stage 3. In general, the increase in cell number followed a similar pattern except that the rise during stage 2 was less pronounced compared to the corresponding volume increase.

**Figure 5.**
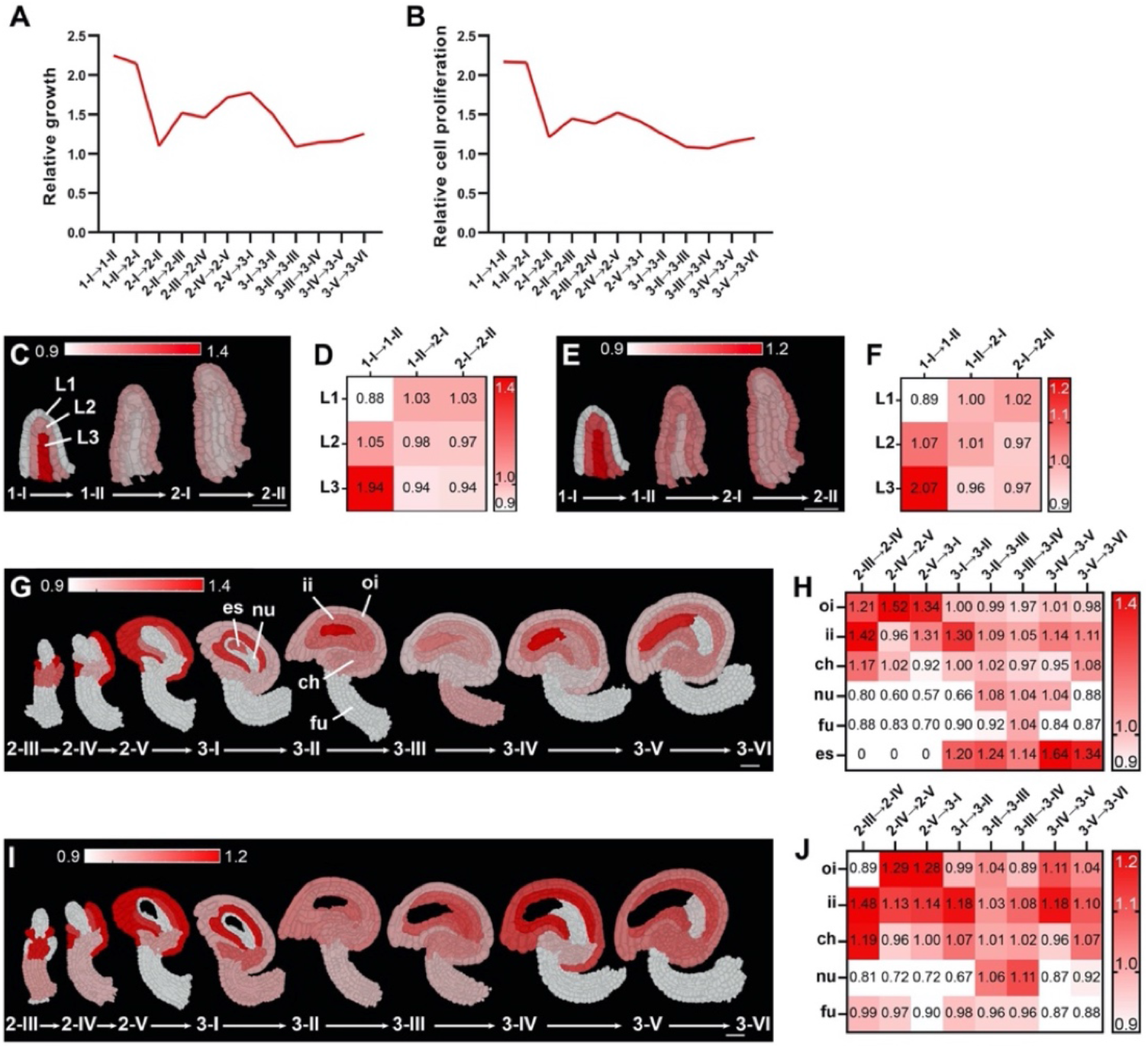
Growth dynamics of Arabidopsis ovule development. (A, B) Plots indicating the overall volume or cell number increase from one stage relative to the preceding stage. (C-J) Optical mid-sections and heat maps depicting relative tissue growth across the different ovule stages. Stages are indicated. Heatmap values indicate ratios. (C, D, G, H) Tissue-specific growth rate. (E, F, I, J) Tissue-specific cell proliferation rate. Abbreviations: ch, chalaza; es, embryo sac; fu, funiculus; ii, inner integument; nu, nucellus; oi, outer integument.

We then investigated the relative growth of the major tissues (Fig. 5C-J). To this end we took the ratio between the mean total volume or mean total cell number of a tissue for two consecutive stages and divided it by the corresponding ovule growth rate. The ratio infers the tissue growth rate with respect to ovule growth. Values above 1 indicate that the tissue is growing at a higher rate than overall ovule growth rate and values below 1 indicate the opposite. We observed that the L3 showed the most relative growth and the L1 the least during stage 1-I (Fig. 5C-F). During stages 1-II to 2-II all three layers contributed similarly to the overall growth. From stage 2-III to 3-VI the nucellus, chalaza, integuments, and eventually the embryo sac showed dynamic changes during development (Fig. 5G-J). For example, up to stage 3-I the outer integument exhibited more relative growth than the inner integument while from stage 3-I on the relative growth pattern was reversed.

In summary, the results reveal ovule growth to be dynamic both in terms of overall growth and the respective relative contributions of individual tissues during development.

### *WUSCHEL* expression is not restricted to the nucellar epidermis

3D digital ovules should allow a detailed investigation of spatial gene expression patterns and with cellular resolution. To test this assumption, we analyzed the expression of *WUSCHEL* (*WUS*, AT2G17950) in the young ovule. *WUS* controls stem cell development in the shoot apical meristem (Gaillochet et al., 2015; Mayer et al., 1998). *WUS* is also active during ovule development where it is expressed in the nucellus and controls germ line development and the organization of the central region (Gross-Hardt et al., 2002; Lieber et al., 2011; Sieber et al., 2004; Tucker et al., 2012; Zhao et al., 2017). Investigation of the *WUS* expression pattern by in situ hybridization or reporter gene analysis left open the question whether *WUS* expression is restricted to the nucellar epidermis at all times or whether it is also present in interior tissue. To address this issue we took advantage of a Col reporter line carrying a reporter for *WUS* promoter activity (pWUS::2xVENUS:NLS::tWUS (pWUS)) (Zhao et al., 2017). We generated a total of 67 3D digital ovules from the pWUS reporter line covering stages 1-I to 2-III (9 ≤ n ≤ 21 per stage).

We first asked when the pWUS signal became detectable during ovule development. We found that with two exceptions pWUS signal could robustly be observed starting with ovules carrying 50 or more cells (Fig. 6A,C). Of the 38 digital ovules spanning stage 1 only two cases with fewer than 50 cells showed either one (1048_A, 32 cells) or two (805_E, 48 cells) cells expressing pWUS signal while all ovules with 50 or more cells showed pWUS signal. The morphological distinction between stages 1-I and 1-II had previously not been clearly defined (Schneitz et al., 1995). Thus, we used pWUS expression as a convenient marker to more precisely discriminate between the end of stage 1-I (ovules with fewer than 50 cells) and the start of stage 1-II (ovules with 50 or more cells).

**Figure 6.**
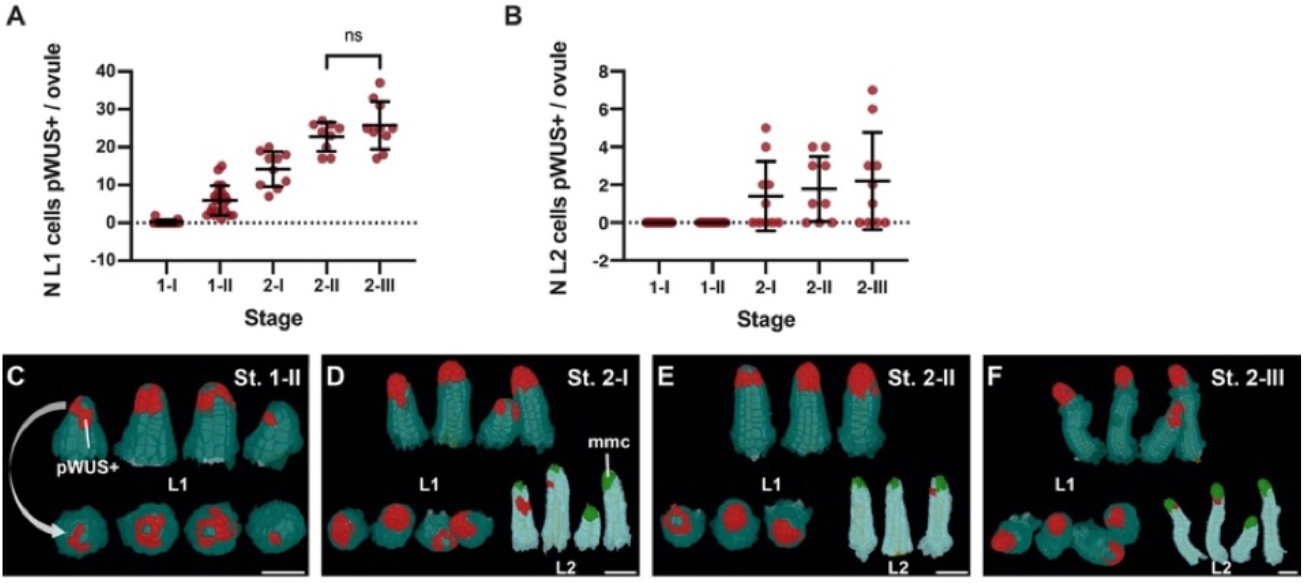
Expression pattern of the pWUS reporter. (A) Plot showing the number of L1 cells per ovule expressing pWUS across stage 1-I to 2-III. (B) Plot showing the number of L2 cells per ovule expressing pWUS across stage 1-I to 2-III. (C-F) 3D cell meshes displaying L1 and L2 cells expressing pWUS in red from stage 1-II to stage 2-III. Data points indicate individual ovules. Mean ± SD are represented as bars. Scale bars: 20 μm.

Next, we explored the elaboration of the spatial pWUS expression pattern during early primordium development (Fig. 6C-F). Reporter signal was initially detected in the epidermis of the distal tip of the primordium with individual cells or in small irregular patches of cells exhibiting reporter expression. The patchiness of the signal continued through stage 2-II. By stage 2-III, however, most of the epidermal cells of the nucellus exhibited pWUS signal. A pWUS reporter signal in the nucellar epidermis at early stage 2 is consistent with previous reports (Tucker et al., 2012; Zhao et al., 2017). However, we noticed that starting with stage 2-I about half of the ovules also exhibited reporter signal in a few cells of the first subepidermal layer (L2) (Fig. 9B, D-F). In those instances, between one to seven L2 cells were found to express the reporter. All the pWUS-expressing L2 cells were neighboring the MMC and resided next to the bottom of the MMC. We did not detect the reporter signal in the MMC itself. Finally, we did not detect signals in the central region or the integuments. Thus, our data indicate a strong temporal control of *WUS* expression with pWUS expression becoming visible in the 50 cell primordium. By contrast, spatial regulation of *WUS* appears less strict as indicated by the variable and spotty expression of pWUS during stages 1-II to 2-I.

### Ovule primordia grow evenly

To gain more detailed insight into specific aspects of ovule development we explored a range of stage and tissue-specific cellular properties of the 3D digital ovules. We first investigated early primordium development up to the appearance of the MMC at the distal tip of the primordium but prior to the initiation of integument development (stages 1-I, 1-II, and 2-I). To this end we generated a dataset comprising 138 digital primordia of stage 2-I or younger. This dataset included ovules from the high-quality dataset described above but also encompassed ovule primordia that did not make the cut for the high-quality dataset as they possessed slightly higher percentages of undersegmented cells (more than four percent but less than nine percent of missegmented cells per primordium). This approach increased sample size and still allowed error-free determination of cell numbers (by including information from the TO-PRO-3 channel). Moreover, we could also determine the total volume of the primordium, by summing up the volumes of its constituting cells, as undersegmenation errors minimally influence this parameter. Finally, this dataset was useful to determine the PD extension of the primordia.

We first wanted to assess if primordium outgrowth occurs evenly or if there is evidence for distinct growth phases. We ordered the individual digital primordia according to increasing PD extension (primordium height), primordium volume, or cell number (Fig. 7A-D). We did not detect distinct subclasses of primordia but observed steady and continuous rises in primordium volume and cell number. Taken together, the data support the notion that from the youngest detectable primordia to late stage 2-I primordia outgrowth does not undergo major fluctuations.

**Figure 7.**
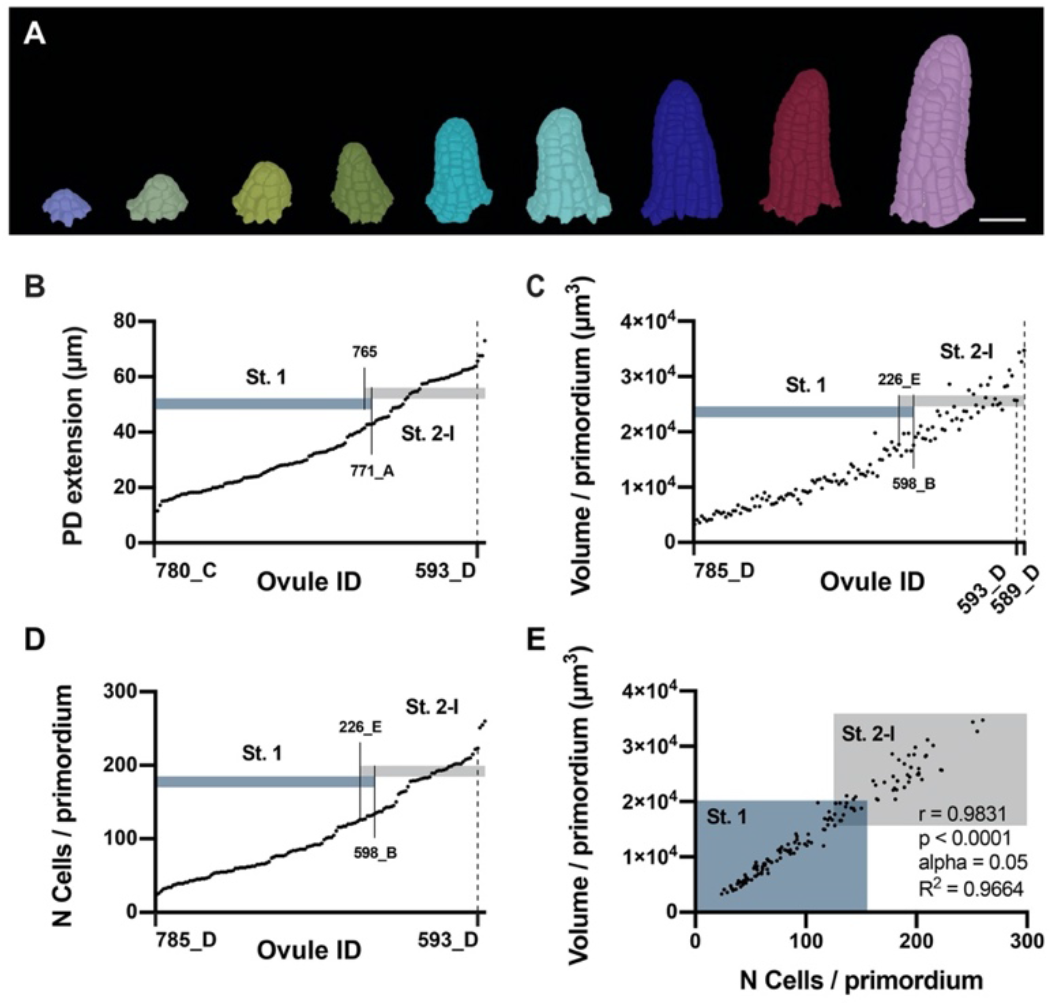
Ovule primordia grow in a continuous fashion. (A) Developmental series of 3D cell meshes of ovule primordia. (B) Plot showing an ordered array of the PD extension of ovule primordia from early initiation to the end of stage 2-I. (C) Plot indicating the total volume of primordia ordered according to increasing volume. (D) Plot depicting the total number of cells in the ovules ordered according to the increasing number of cells. (E) Plot representing the correlation between total number of cells and total volume per primordia. Data points indicate individual ovules. Scale bar: 20 μm.

Next we assessed how cell number per primordium and total primordium volume relate to each other. We observed a positive correlation (Fig. 7E). Interestingly, however, data points representing stage 2-I primordia showed noticeably more scattering than the data points of stage 1 primordia. This indicates that for stage 2-I primordia, the total volume in relation to cell number is more variable than for stage 1 primordia. One can also detect more variation in total volume per primordium in the ordered list of total volume per primordium (Fig. 7C). These results are consistent with the notion that growth of stage 1 primordia is under comparably tighter growth control while growth during stage 2-I ovules is more variable (see also below).

We then addressed the question if there is a rapid transition between stage 1-II and stage 2-I. Stage 2-I is defined by the emergence of the large L2-derived MMC at the distal tip of the primordium (Schneitz et al., 1995). To this end we determined which stage 1 ovules exhibited maximum values for the three parameters mentioned above and which stage 2-I ovules featured the smallest values, respectively (Fig. 7B-D). Taking these considerations into account, ovule primordia grow to a volume of about 1.8 × 10^4^ μm^3^, or a cell number range of approximately 125 to 135 total cells, and to a height range of about 41 to 43 μm when they enter stage 2-I. For each of the three parameters we noticed a small number of ovules that fell into the range of overlap: five for the PD extension and seven each for total volume of primordium and total cell number per primordium. These numbers account for 3.6 percent and 5.1 percent of the total of 138 scored ovules, respectively. This result indicates that ovule primordia grow to a certain size and then rapidly transition from stage 1-II to stage 2-I.

Assessment of the cell volume of the L2 cells in our datasets of stage 1-II and stage 2-I digital ovules combined with visual inspection of the digital ovule primordia revealed that the MMC can easily be distinguished based on its presence in the L2 and its comparably large cell volume (Fig. 8A,B). At stage 2-I we measured an average MMC volume of 543 μm^3^ (543.3 ± 120.6). The smallest MMC at stage 2-I possessed a volume of 339.2 μm^3^, well within the range observed previously (Lora et al., 2017). The largest L2 cell at stage 1-II had a volume of 297.6 μm^3^. We observed a mean cell volume of stage 1-II L2 cells of 153.5 μm^3^ ± 48.3 (n = 335) and the volume of this large cell was well beyond the 75% percentile (187.7 μm^3^). However, visual inspection revealed that this cell did not reside at the tip of the primordium while all the L2 cells located at the tip of the different stage 1-II primordia showed a cell volume that was close to the mean value. While we cannot exclude that MMC development starts already earlier than stage 2-I we propose that by definition a stage 2-I MMC has a minimal cell volume of 335 μm^3^.

**Figure 8.**
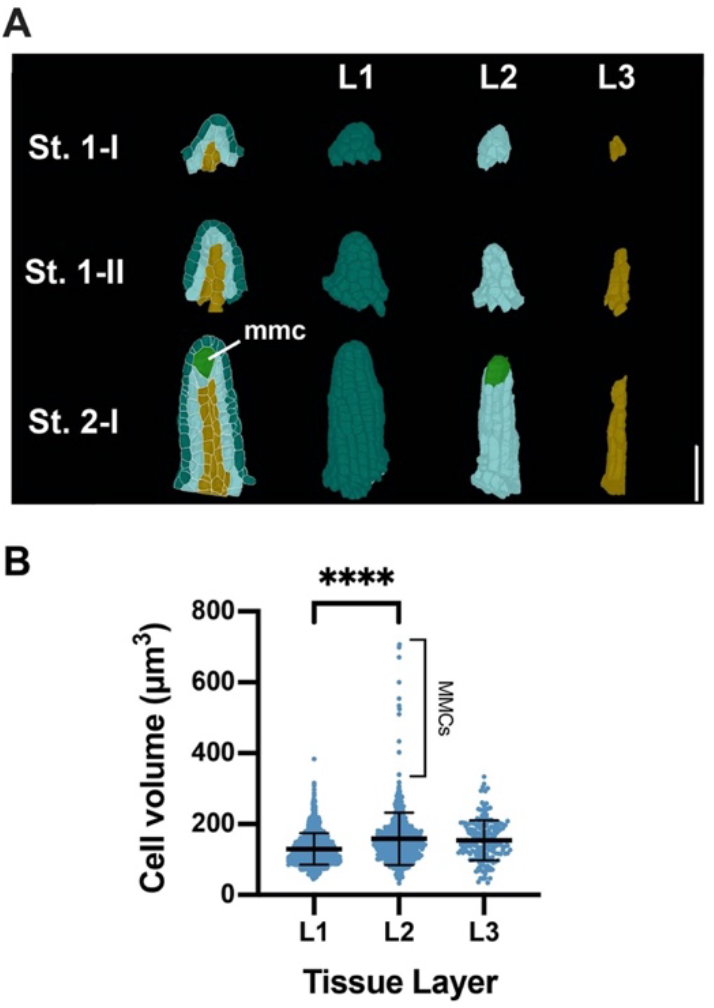
Radial tissue organization in ovule primordia. (A) Mid-sagittal section view of ovule primordia and extracted 3D cell meshes of L1, L2, L3 layers separately viewed in 3D from stage 1-I to 2-I. (B) Plot depicting the cell volumes of the L1, L2, L3 per ovule at stage 2-I. Data points indicate the volumes of individual cells. Mean ± SD are represented as bars. Asterisks represent statistical significance (****, P<0.0001; Student’s t test). Abbreviations: mmc, megaspore mother cell. Scale bar: 20 μm.

### Ovule primordium slanting indicates early onset of gynapical-gynbasal polarity

Upon inspecting the 3D meshes of pistil fragments we noticed that many ovule primordia were positioned at a hitherto undescribed slant relative to the placenta surface (Fig. 9). We quantified the slant by measuring the PD distances of the shorter and longer sides of primordia of different stages. We observed that the slant was barely noticeable at stage 1-I, became more tangible during stage 1-II, and was prominent by stage 2-I. At stage 2-III initiation of outer integument occurred more on the side of the small angle of the slant indicating that the small angle was positioned on what will become the gynbasal side of the developing ovule. Thus, we propose that the gynapical-gynbasal polarity of the ovule becomes morphologically detectable already during stage 1 and not only later at stage 2-III when outer integument initiation is discernible at the gynbasal side of the primordium.

**Figure 9.**
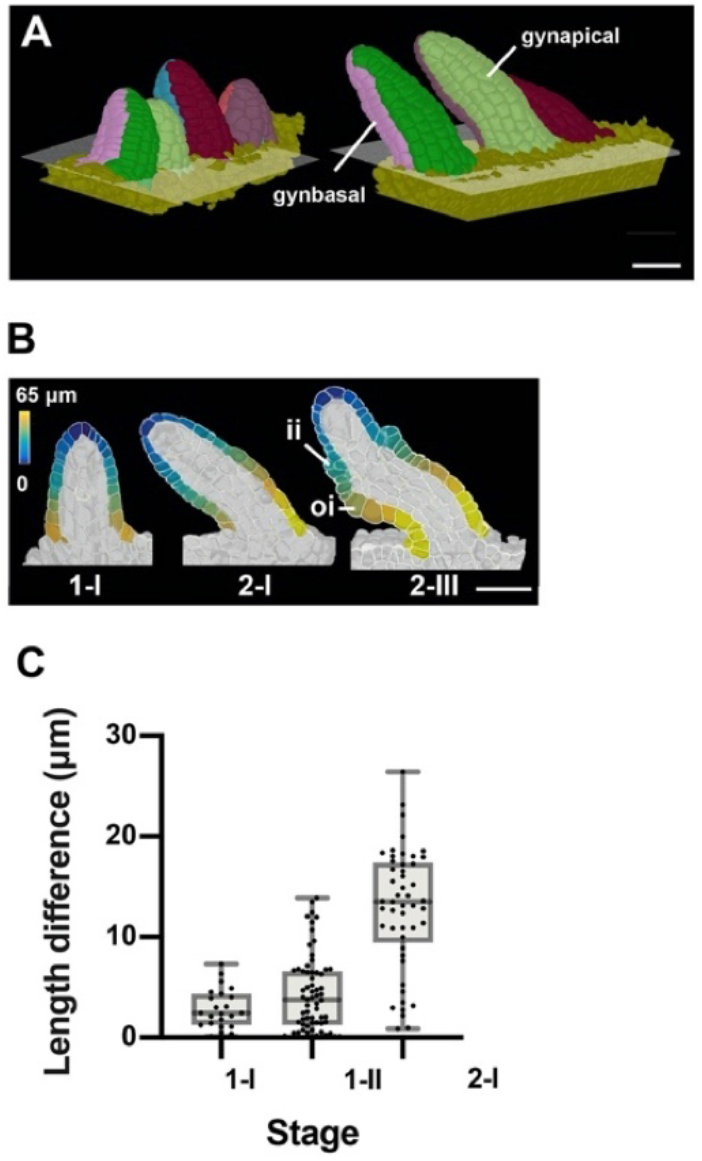
Ovule primordia slanting. (A) 3D mesh view with multiple ovules from the same carpel showing unslanted ovules at stage 1-I and slanted ovules at stage 2-I. The 3D grid represents the surface of the placenta. Color labels depict the gynapical or gynbasal cells, respectively. (B) 2D section view of a 3D cell mesh from an early stage to 2-III ovule. The heatmap on the epidermal cells of gynapical and gynbasal halfs depicts the quantified distance value between individual measured cells to the distal tip of primordia. (C) Plot depicting the extent of slanting, quantified by the difference in maximal length on the gynapical and gynbasal sides of ovule at stages 1-I, 1-II and 2-I. Data points indicate individual ovules. Mean ± SD are represented as bars. Scale bars: 20 μm.

### A few scattered asymmetric cell divisions initiate a parenchymatic inner integument layer

A new cell layer is produced by the inner integument shortly before fertilization (Debeaujon et al., 2003; Schneitz et al., 1995). The ii1 or endothelial layer originates this additional cell layer (ii1’) by periclinally-oriented asymmetric cell divisions (Fig. 10A). Cells of the ii1’ layer are immediately distinguishable from ii1 cells by their altered shape and staining characteristics (Schneitz et al., 1995). Moreover, they do not express a transcriptional reporter for the epidermis-specific *ARABIDOPSIS THALIANA MERISTEM L1* (*ATML1*) gene (Debeaujon et al., 2003). Finally, during early seed development, ii1 cells will produce tannins while ii1’ cells will contribute to parenchymatic ground tissue. Thus, cambium-like activity of the ii1 layer results in asymmetric cell divisions, thickening of the inner integument, and formation of distinct tissues with separate functions.

**Figure 10.**
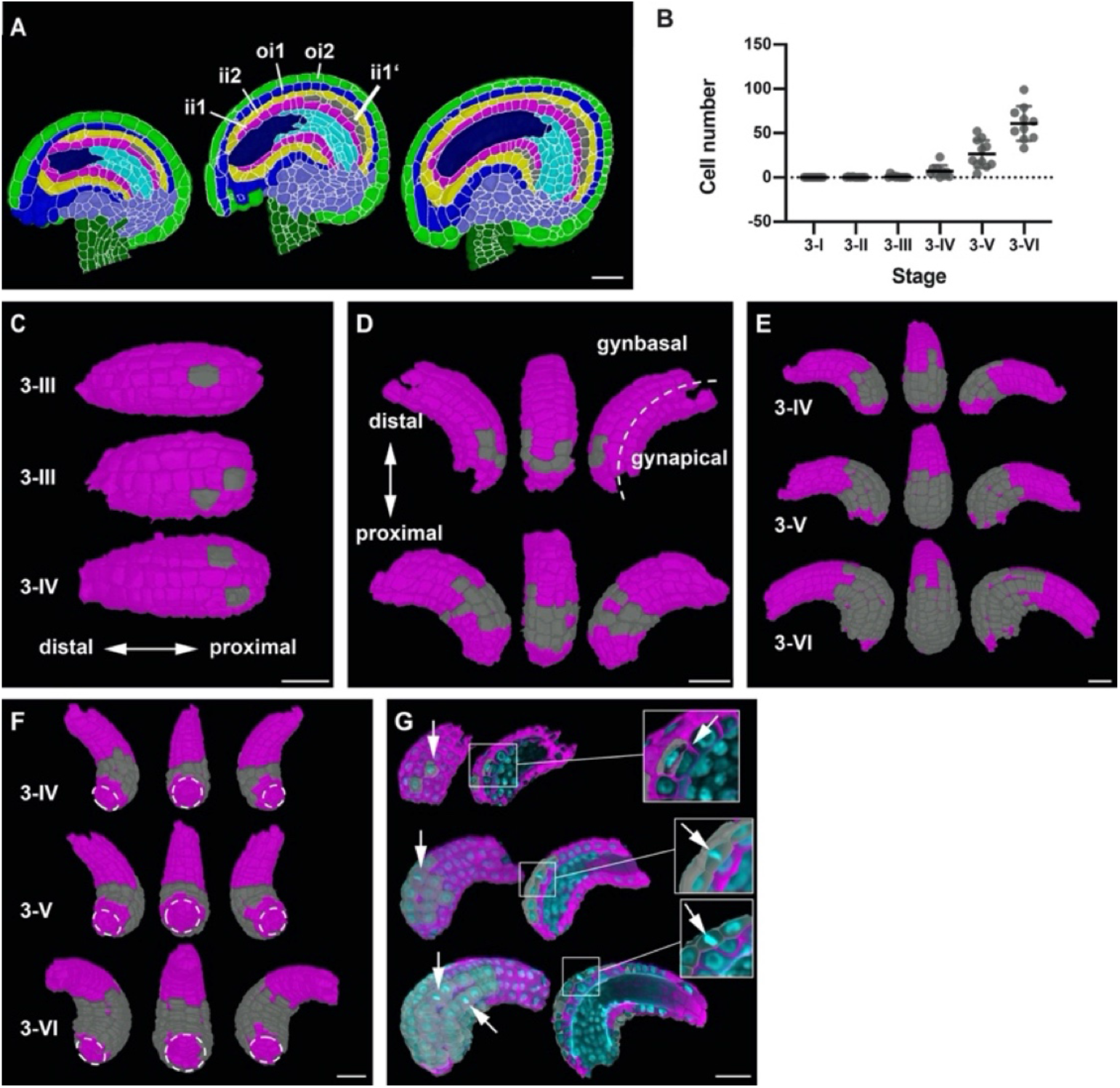
Formation of the parenchymatic inner integument layer (ii1’). (A) Mid-sagittal section of wild-type ovule at stages 3-IV, 3-V and 3-VI, showing the initiation of a new cell layer (ii1’) in the adaxial inner integument (endothelium/ii1). (B) Plot depicting the number of cells of the developing ii1’ layer. Data points indicate individual ovules. Mean ± SD are represented as bars. (C) 3D top surface view of the adaxial inner integument at stages 3-III and 3-IV showing the occurrence of the first few pioneer cells of the ii1’ layer. (D) 3D side surface view of adaxial inner integument depicting the pattern of occurrence of ii1’ cells. (E) 3D side surface view of adaxial inner integument at later stages of 3-IV, 3-V and 3-VI where the emergent tissue layer is observed to be a patch of connected cells present only at the proximal region of the inner integument. (F) 3D bottom surface view of ii1’ layer at different stages highlighting the formation of a ring-like structure of connected cells covering the proximal half of the inner integument. (G) Section view of the 3D cell meshes of adaxial inner integument with the overlaid nuclei z-stack displaying a periclinal division (top) and anticlinal divisions (center, bottom) in the ii1’ layer. Abbreviations: ii1, adaxial inner integument (endothelium); ii1’, parenchymatic inner integument layer; ii2, abaxial inner integument; oi1, adaxial outer integument; oi2, abaxial outer integument. Scale bars: 20 μm.

The cellular basis and 3D architecture of ii1’ layer formation is poorly understood. We followed its formation through all of stage 3 with the help of our 3D digital ovules (Fig. 10B-G). To this end we removed the outer integument and layer ii2 in MGX. We could observe the ii1’ layer at various stages of development in 36 digital ovules. In contrast to what was previously described we already observed first signs of ii1’ formation at stage 3-II (Fig. 10B,C). Two out of the ten digital ovules showed one cell of this layer. At stage 3-III 4/11 and at stage 3-VI 9/10 digital ovules showed at least one ii1’ layer cell, respectively. By stages 3-V and 3-VI all digital ovules exhibited this layer. The number of ii1’ layer cells increased to 60.8 ± 19.5 at stage 3-VI. Thus, at this stage the average ii1’ layer consisted of 13.4 percent of the total cells of the inner integument and with a volume of 1.53 × 10^4^ μm^3^ ± 0.5 × 10^4^ μm^3^ contributed 14.4 percent to its total volume.

With respect to the spatial organization of the ii1’ layer we observed single cells or patches of cells that were located on both lateral sides of the gynbasal inner integument at stages 3-II and 3-III (Fig. 10C). Due to further divisions a patch of connected cells became visible at later stages. Proximal-distal and lateral extension of the ii1’ layer occurred until it showed a ring-like structure covering the proximal half of the inner integument at stage 3-VI (Fig. 10D-F). The boundary of the ii1’ layer was not smooth but exhibited an irregular appearance.

We then asked if the ii1’ layer is generated entirely through periclinal cell divisions of ii1 cells or if anticlinal cell divisions in existing ii1’ cells contribute to the formation of this layer. To this end we scored 978 cells of the ii1’ layers across the 36 ovules exhibiting a ii1’ layer and identified 14 cells in mitosis as indicated by the presence of mitotic figures in the TO-PRO-3 channel (Fig. 10G). Only one of the 14 cells showed a periclinal cell division that generated a cell of the ii1’ layer. Interestingly, however, the other 13 mitotic cells were experiencing anticlinal cell divisions.

Taken together, the results suggest that initiation of the ii1’ layer does not involve simultaneous asymmetric cell divisions in many ii1 cells resulting in the formation of a ring or large patch of ii1’ tissue. Rather, asymmetric periclinal cell divisions occur only in a few spatially scattered founder cells distributed within the ii1 cell layer. Further anticlinal cell divisions in the ii1’ daughters of the founder cells then result in the formation of a continuous ii1’ cell layer with irregular edges.

### *INNER NO OUTER* affects development of the nucellus

The YABBY transcription factor gene *INNER NO OUTER* (*INO*, AT1G23420) is an essential regulator of early pattern formation in the ovule (Baker et al., 1997; Balasubramanian and Schneitz, 2002, 2000; Meister et al., 2002; Schneitz et al., 1997; Sieber et al., 2004; Villanueva et al., 1999). Plants with a defect in *INO* carry ovules that fail to form an outer integument. Moreover, development of the female gametophyte is usually blocked at the mono-nuclear embryo sac stage. In situ hybridization and reporter gene studies revealed that *INO* is exclusively expressed in the abaxial layer of the outer integument (oi2). Although the ovule phenotype of *ino* mutants has been well characterized it remained unclear if the inner integument and the nucellus, apart from the defect in embryo sac development, are affected in *ino* mutants. To address these and other issues (see below) we generated *ino-5*, a putative null allele of *INO* in Col-0 that was induced by a CRISPR/Cas9-based approach (see Materials and Methods) and performed a quantitative phenotypic analysis of *ino-5* ovules using a dataset of 119 3D digital *ino-5* ovules covering stages 1-II to 3-VI (3 ≤ n ≤ 20 per stage) (Fig. 11A). Ovules lacking *INO* activity are difficult to stage as they lack many of the distinct criteria that define the different stages of wild-type ovule development. To circumvent this problem we staged *ino-5* ovules by comparing the total number of cells and the total volume of ino-5 3D digital ovules to the corresponding stage-specific values of wild-type 3D digital ovules for which the outer integument had been removed (Table S2).

**Figure 11.**
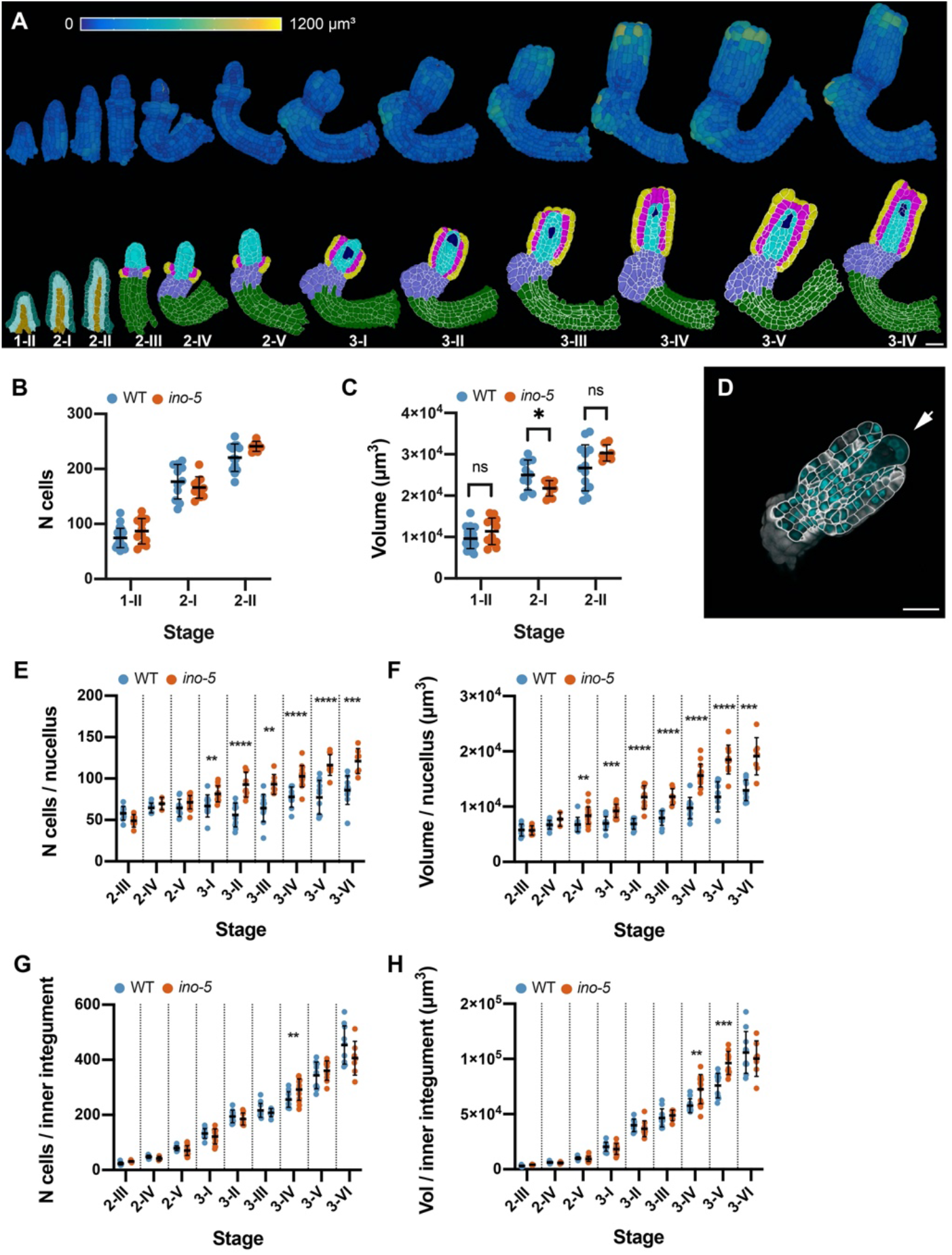
Quantitative analysis of the *ino-5* digital ovule atlas. (A) Surface view of 3D cell meshes showing heatmaps of cell volumes of *ino-5* ovules from early to late stage of development. The bottom row shows 3D mid-sagittal section views depicting the cell type organization in *ino-5* mutant ovules. (B) Plot showing the total number of cells in wild-type and *ino-5* at early stages of ovule development. (C) Plot showing the total volume of wild-type and *ino-5* ovules at early stages of ovule development. Section view of the cell boundary z-stack (white) overlayed with the z-stack of stained nuclei showing a four-nuclear embryo sac in an ino-5 ovule. (E, F) Plots comparing total cell number and tissue volume of the nucellus between wild type and ino-5 at different stages. (G, H) Plots comparing total cell number and tissue volume of the inner integument between wild type and ino-5 at different stages. Data points indicate individual ovules. Mean ± SD are represented as bars. Asterisks represent statistical significance (ns, P ≥ 0.5; *, P < 0.05; **, P < 0.01, ***, P < 0.001; ****, P < 0.0001; Student’s t test). Scale bars: 20 μm.

We first analyzed the total cell number per primordium and the total volume per primordium for *ino-5* ovules of stages 1-II to 2-II. We did not find robust differences between *ino-5* and wild type for the two parameters (Fig. 11B,C). This was to be expected as *INO* expression became first detectable around stage 2-II/III (Balasubramanian and Schneitz, 2000; Meister et al., 2004, 2002; Villanueva et al., 1999). We then asked if *INO* affects the development of the nucellus and the inner integument. To this end we analyzed cellular features of the two tissues of *ino-5* for stages 2-III to 3-VI. We first investigated nucellar development. As a rule, and as previously described, we observed that embryo sac development did not extend beyond the mono-nuclear embryo sac stage although up to four-nuclear embryo sacs could be detected as well (Fig. 11D). We then assessed cell number and tissue volume of the nucellus of *ino-5* (excluding the embryo sac). We observed an elevated number of nucellar cells in *ino-5* ovules starting from late stage 2-V/stage 3-I. Wild-type stage 3-I ovules featured 66.9 ± 13.4 cells per nucellus while *ino-5* carried 81.41 ± 9.6 cells per nucellus (Fig. 11E) (Table 3). The nucellar volume of *ino-5* (0.9 ± 0.1 μm^3^) was also increased relative to wild type (0.7 ± 0.1 μm^3^) (Fig. 11F). The results indicate that *INO* not only affects ontogenesis of the embryo sac but also restricts cell number and nucellus volume during nucellus development.

**Table 3.** Cell numbers and total volumes of different tissues in *ino-5*.

|  | Tissue |  |  |  |  |  |  |  |
| --- | --- | --- | --- | --- | --- | --- | --- | --- |
| Stage<sup>a</sup> | Nucellus |  | Inner Integument |  | Funiculus |  | Central region |  |
|  | N cells | Volume<br>(x10<sup>4</sup> μm<sup>3</sup>) | N cells | Volume<br>(x10<sup>4</sup> μm<sup>3</sup>) | N cells | Volume<br>(x10<sup>4</sup> μm<sup>3</sup>) | N cells | Volume<br>(x10<sup>4</sup> μm<sup>3</sup>) |
| 2-III | 49 ± 7.6 | 0.5 ± 0.08 | 31.3 ± 3.2 | 0.4 ± 0.03 | 189.0 ± 29 | 2.4 ± 0.4 | 35.8 ± 37.1 | 0.3 ± 0.3 |
| 2-IV | 69.7 ± 7.5 | 0.7 ± 0.1 | 41.6 ± 8.3 | 0.5 ± 0.1 | 202.7 ± 25 | 2.2 ± 0.4 | 68.3 ± 6.6 | 0.6 ± 0.2 |
| 2-V | 71.4 ± 8.1 | 0.8 ± 0.1 | 70.7 ± 17.7 | 0.9 ± 0.2 | 286.4 ± 35.2 | 3.4 ± 0.5 | 108.1 ± 19.3 | 1.2 ± 0.3 |
| 3-I | 81.4 ± 9.5 | 0.9 ± 0.1 | 122.0 ± 27.3 | 1.8 ± 0.5 | 379.0 ± 29.7 | 4.7 ± 0.5 | 134.0 ± 23.1 | 1.8 ± 0.4 |
| 3-II | 92.6 ± 15.0 | 1.1 ± 0.2 | 185.1 ± 21.9 | 3.9 ± 0.7 | 402.9 ± 32.6 | 5.1 ± 0.7 | 176.9 ± 24.6 | 2.6 ± 0.4 |
| 3-III | 93.0 ± 12.0 | 1.1 ± 0.1 | 208.4 ± 14.6 | 4.8 ± 0.5 | 451.6 ± 32.7 | 5.9 ± 0.7 | 202.9 ± 24.1 | 3.2 ± 0.5 |
| 3-IV | 102.7 ± 12.8 | 1.5 ± 0.2 | 296.9 ± 43.5 | 7.3 ± 1.3 | 481.3 ± 45.7 | 6.7 ± 0.5 | 215.4 ± 30.7 | 3.6 ± 0.7 |
| 3-V | 116.3 ± 12.5 | 1.8 ± 0.2 | 360.8 ± 35.2 | 9.6 ± 1.0 | 522.6 ± 47.4 | 7.2 ± 1.1 | 239.6 ± 31.2 | 4.2 ± 0.7 |
| 3-VI | 121.1 ± 15.2 | 1.9 ± 0.3 | 406.1 ± 61.3 | 10.0 ± 1.6 | 599.6 ± 40.6 | 8.8 ± 0.8 | 254.4 ± 48.5 | 4.5 ± 0.9 |

<sup>a</sup>Number of 3D digital ovules scored: 3 (stages 2-IV), 6 (stages 2-III), 7 (stages 3-III, 3-VI), 9 (stages 3-II), 10 (stages 3-V), 14 (stages 3-I, 3-IV), 20 (stages 2-V). Values represent mean ± SD.

We then investigated cell number and tissue volume for the inner integument of *ino-5* (Fig. 11G,H) (Tables 2 and 3). We found that the two parameters did not deviate noticeably from wild type except for stages 3-IV and 3-V where we observed a small but perceivable increase in cell number and tissue volume. However, this increase appeared to be transient as both parameters were normal in *ino-5* ovules of stage 3-VI. These findings indicate that *INO* does not exert a major influence on the monitored cellular characteristics of the inner integument.

### Ovule curvature represents a multi-step process

The curvature of the Arabidopsis ovule constitutes an interesting and unique morphogenetic process. It raises the question if curvature can be subdivided into distinct, genetically controlled steps. During stages 2-III to 2-V of wild-type ovule development a prominent kink forms in the ovule primordium resulting in the nucellus pointing towards the gynapical side of the ovule (Figures 1A, 11A and 12A). This early gynapically-oriented kink was absent in *ino-5* ovules (Fig. 12A). In later stage *ino-5* ovules, we found aberrant growth on the gynapical side of the chalaza and the nucellus pointing more to the gynbasal side of the ovule (Fig. 12A). Apart from the absence of the regularly oriented kink in *ino* ovules there is an obvious effect on the curvature of the ovule since the nucellus and inner integument develop into straight rather than curved structures.

**Figure 12.**
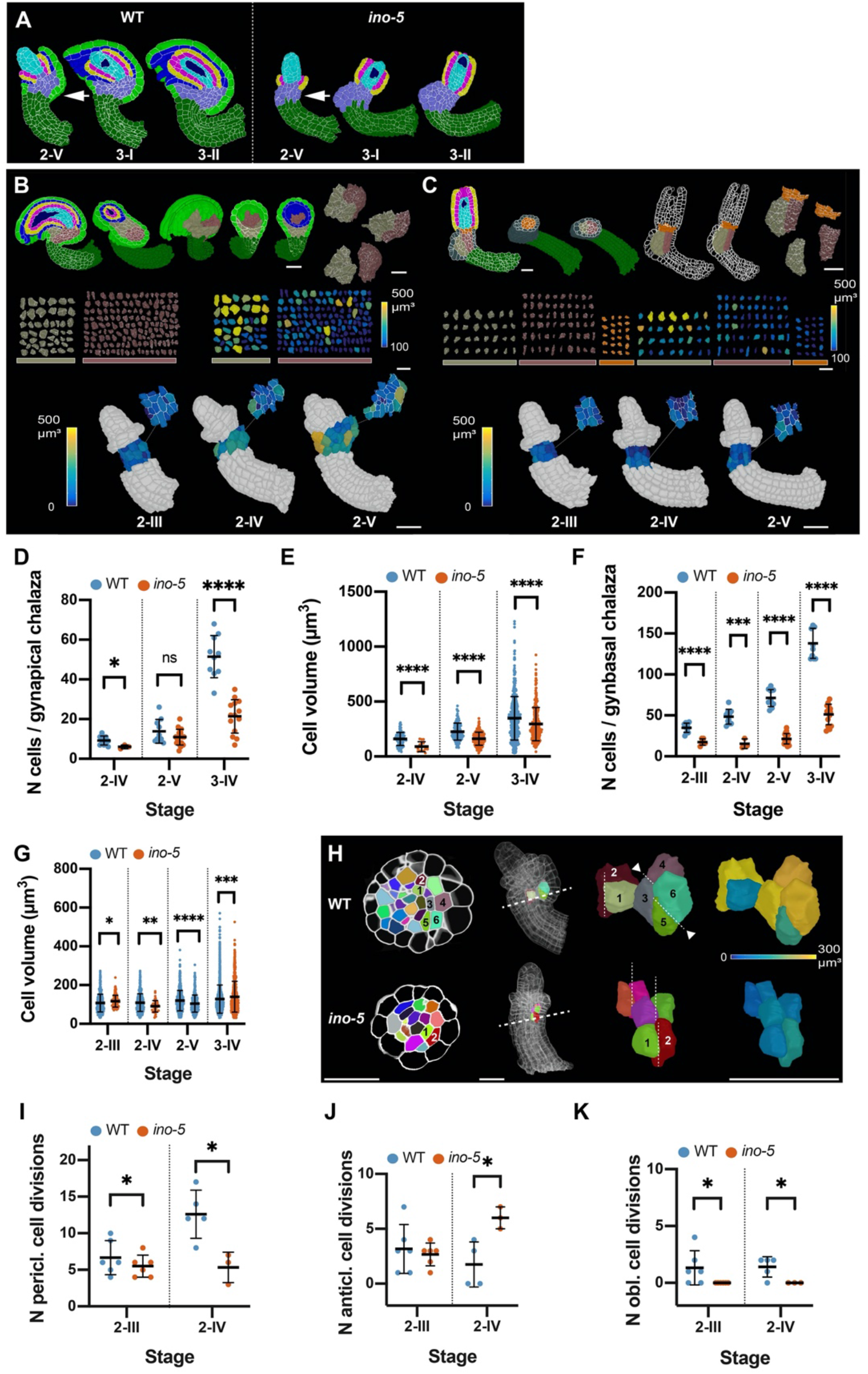
Growth patterns forming the subepidermal central region in wild-type and *ino-5* ovules. (A) Mid-sagittal section view of the cell-type-labelled 3D cell meshes of wild type and *ino-5* showing the differences in tissue organization across stage 2-V to 3-II. (B) Representation of the two chalazal regions in mature wild-type ovules from different perspectives, shaving off cell layers and extracting out individual cells in 3D to visualize cell morphology and cell volume. The bottom row shows the 3D cell meshes of wild-type ovules from stage 2-III to 2-V. The outer integument has been removed to reveal cell volumes of the two subepidermal regions in wild type. The small figure on the top right represents the section view of the 3D cell meshes of the corresponding ovule. (C) Mature *ino-5* ovule depicting the chalazal region and extracted cells types in 3D similar to (B). The bottom row shows 3D cell meshes of *ino-5* ovules from stage 2-III to 2-V. The epidermis has been removed to reveal the cell volumes of the two subepidermal regions in *ino-5*. The small figure on the top right represents the section view of the 3D cell meshes of the corresponding ovule. (D) Comparison of cell numbers between wild type and *ino-5* in the gynapical chalaza of stage 2-IV, 2-V and 3-IV ovules. Data points indicate individual ovules. (E) Comparison of cell volume between wild type and *ino-5* in the gynapical chalaza of stage 2-IV, 2-V and 3-IV ovules. Data points indicate volume of individual cells. (F) Plot comparing the cell number in the gynbasal chalaza between stage 2-IV, 2-V, and 3-IV wild-type and *ino-5* ovules. Data points indicate individual ovules. (G) Plot comparing the cell volume in the gynbasal chalaza between stages 2-IV, 2-V and 3-IV wild-type and *ino-5* ovules. Data points indicate volumes of individual cells. (H) Left section: radial section view depicting the division patterns observed in the chalazal region in wild-type and *ino-5* ovules. The dashed line indicates the section plane shown on the left. Right section: Oblique 3D view of the cells numbered in the radial section view. Dashed lines indicate the cell division plane. Arrowheads highlight oblique periclinal cell division planes. The heat map indicates the volume. (I) Plot comparing the number of periclinal divisions in wild-type and *ino-5* ovules. (J) Plot comparing the number of longitudinal-anticlinal cell divisions in wild type and *ino-5*. (K) Plot comparing the number of oblique divisions in wild type and *ino-5*. Data points indicate individual cell division. Mean ± SD are represented as bars. Asterisks represent statistical significance (ns, P ≥ 0.5; *, P < 0.05; **, P < 0.01; ***, P < 0.001; ****, P < 0.0001; Student’s t test). Scale bars: 20 μm.

### Two morphologically distinct subepidermal regions contribute to kink formation in the proximal chalaza

Formation of the kink in the early ovule indicates that differential growth patterns in the central region may underlie this process. The differences in kink formation between wild-type and *ino-5* ovules prompted us to investigate the cellular patterns of the subepidermal central region of the two genotypes in more detail. Previous genetic results as well as evolutionary considerations implied that the central region or chalaza can be subdivided into distal and proximal tiers flanked by the inner and outer integuments, respectively (Baker et al., 1997; Endress, 2011; Gasser and Skinner, 2019; Sieber et al., 2004). We focussed on the cellular architecture of the proximal chalaza which constitutes the majority of the central region. In wild-type ovules, we observed that the outer integument was to a large extent made of epidermis-derived cells. However, we also identified two groups of subepidermal cells in the proximal chalaza that contribute to the development of the central region in a differential manner with one group of cells also making an obvious subepidermal contribution to the outer integument (Fig. 12B). Due to their locations the two groups of cells were termed gynapical and gynbasal chalaza, respectively. By stage 3-IV the two cell populations were easily distinguished based on cell shape, volume, and number of the containing cells. We observed fewer but bigger cells in the gynapical chalaza and the opposite for the gynbasal chalaza (Fig. 12B, D-G). Furthermore, we noticed that the two tissues eventually exhibit distinct shapes with the gynapical chalaza showing two “wing-like” lateral extensions while the gynbasal chalaza acquires a radially symmetric, rod-like shape (Fig. 12B).

In stage 2-III/2-IV wild-type ovules most cells of the proximal chalaza appeared relatively homogeneous in size and shape (Fig. 12B,D-G) (Table 4). At stage 2-III, we counted 6.7 ± 2.3 periclinal cell division planes in the subepidermal proximal chalaza (Fig. 12H,I). This number increased to 12.6 ± 3.3 by stage 2-IV. We also observed a small number of division planes oriented in a longitudinal-anticlinal fashion (3.2. ± 2.2 at stage 2-III and 1.8. ± 2.1 at stage 2-IV) (Fig. 12H,J). These division planes were distributed throughout the subepidermal proximal chalaza. Interestingly, we also found a few oblique division planes (1.3 ± 1.5 at stage 2-III and 1.4 ± 0.9 at stage 2-IV) (Fig. 12K). They were preferentially found at the gynbasal side of the chalaza just underlying the initiating outer integument (Fig. 12B,H,K). Moreover, we observed that the progeny of the gynbasal oblique cell divisions had undergone asymmetric enlargement with the noticeably bigger cell being located directly adjacent to the large epidermal cells associated with the outer integument initiation (Fig. 12H). We should note that by stage 2-IV a few oblique division planes were also observed at the gynapical chalaza. We propose that the periclinal and longitudinal anticlinal divisions result in a widening of the chalaza. The stage 2-III oblique cell divisions accompanied by asymmetric cell enlargement that are preferentially located at the gynbasal side contribute to early kink formation on the gynbasal side of the primordium.

**Table 4.** Cellular parameters of the proximal chalaza.

|  | Tissue |  |  |  |  |  |
| --- | --- | --- | --- | --- | --- | --- |
| Stage<sup>a</sup> | Gynapical proximal chalaza |  |  | Gynbasal proximal chalaza |  |  |
|  | N cells | Cell volume (μm<sup>3</sup>) | Tissue volume (x10<sup>4</sup> μm<sup>3</sup>) | N cells | Cell volume (μm<sup>3</sup>) | Tissue volume (x10<sup>4</sup> μm<sup>3</sup>) |
| 2-III | - | - | - | 35 ± 5 | 113 ± 41.2 | 0.4 ± 0.08 |
| 2-IV | 9 ± 2 | 160 ± 59.6 | 0.14 ± 0.03 | 48 ± 9 | 109 ± 45.6 | 0.52 ± 0.1 |
| 2-V | 13 ± 6 | 226.4 ± 77.3 | 0.31 ± 0.1 | 71.2 ± 10 | 120 ± 52.7 | 0.85 ± 0.1 |
| 3-I | 32 ± 8 | 270.4 ± 121.3 | 0.87 ± 0.2 | 86 ± 20 | 120 ± 59.5 | 1 ± 0.2 |
| 3-II | 50 ± 8 | 298 ± 156 | 1.5 ± 0.2 | 108 ± 19 | 127 ± 70 | 1.3 ± 0.2 |
| 3-III | 51 ± 13 | 318.1 ± 192 | 1.6 ± 0.35 | 121 ± 32 | 124 ± 68 | 1.5 ± 0.3 |
| 3-IV | 52 ± 11 | 347.5 ± 198.2 | 1.8 ± 0.36 | 138 ± 18 | 128 ± 73 | 1.7 ± 0.2 |
| 3-V | 60 ± 14 | 336.8 ± 227.4 | 2 ± 0.4 | 148 ± 27 | 126 ± 70.5 | 1.8 ± 0.4 |
| 3-VI | 59 ± 6 | 428.7 ± 284.4 | 2.5 ± 0.4 | 209 ± 31 | 132 ± 90.7 | 2.7 ± 0.4 |

<sup>a</sup>Number of 3D digital ovules scored: 10 (stages 2-II- 3-II, 3-IV, 3-VI), 11 (stages 3-III, 3-V).
Values represent mean ± SD

By stage 2-V cells of the gynapical and gynbasal chalaza became more clearly distinguishable (Fig. 13B) (Table 4). At stage 2-V we observed 13 ± 6 cells in the gynapical chalaza with an average cell volume of 226.4 μm^3^ ± 77.3 μm^3^. By stage 3-VI cell number increased to 59 ± 6 cells (a 4.5-fold increase) and the average cell volume to 428.7 μm^3^ ± 284.4 μm^3^ (a 1.9-fold increase). Thus, apart from the gain in cell number the increasing cell volumes of the cells contributed noticeably to the overall volume of the gynapical chalaza. Regarding the gynbasal chalaza we counted 71.2 ± 10 cells in this domain at stage 2-V. At stage 3-VI we found a three-fold increase in cell number with 209 ± 31.5 cells. Average cell volume was 120 μm^3^ ± 52.7 μm^3^ at stage 2-V a value that did not vary much from the cell volume measured at stage 3-VI (Table 4). Thus, the increase in the volume of the gynbasal chalaza appears to be largely driven by cell proliferation. Interestingly, the two tissues reached a similar volume at stage 3-VI.

Development of the gynapical chalaza eventually resulted in a prominent bulging of the outer integument with cells intercalating between the two epidermis-derived layers of the outer integument (Fig. 12B). We never observed gynbasal chalazal cells to intercalate between the two epidermis-derived cell layers of the outer integument indicating that cells of the gynbasal chalaza do not contribute to the development of the outer integument. Thus, the roughly gynapical and gynbasal halves of the outer integument also differ with respect to a subepidermal contribution to their development with subepidermal cells contributing to the gynapical but not the gynbasal outer integument, respectively.

In *ino-5* we observed a different cellular pattern in the proximal chalaza (Fig. 12C-K). We detected an overall smaller number of cells in this tissue and cells tended to not reach the extreme sizes observed in wild type. We noticed overall fewer periclinal division planes in stage 2-III/IV *ino-5* ovules (Fig. 12I). Furthermore, we did not observe oblique division planes (Fig 12H,K). Interestingly, however, the number of longitudinal-anticlinal division planes was enhanced (Fig. 13H,J). Since *INO* expression is restricted to the epidermis and eventually the outer layer of the outer integument (Balasubramanian and Schneitz, 2000; Meister et al., 2004, 2002; Villanueva et al., 1999) (Fig. 10A,B) we propose that *INO* regulates kink formation in a non-cell-autonomous fashion. In the absence of *INO* function, fewer periclinal cell divisions take place. Moreover, symmetric longitudinal-anticlinal cell divisions occur in place of the asymmetric oblique cell divisions in the subepidermal proximal chalaza leading to the absence of the enlarged cells abutting the gynbasal epidermis and a failure of kink formation.

Little cell division and cell enlargement occurred on the gynbasal chalaza of *ino-5* ovules. By contrast, cell divisions and cells with increased size could still be observed at the gynapical side. Eventually, however, the tissue did not extend laterally and thus failed to form the two “wing-like” flaps characteristic for the wild-type gynapical chalaza. As a likely result the tissue formed an obvious frontal protuberance. In addition, the nucellus eventually pointed to the gynbasal side. We assume that the preferential gynapical growth and the absence of the outer integument together lead to the misorientation of the nucellus in *ino-5*.

In summary, the comparison of wild type and *ino-5* ovules indicates that the central region is of complex composition. On the basis of morphological and genetic results, the data suggest that several distinct groups of cells contribute to its interior cellular architecture. The gynapcial and gynbasal chalaza make up most of the subepidermal central region. They eventually reach comparable volumes. The gynapical chalaza shows two “wing-like” lateral extensions while the gynbasal chalaza exhibits a radially symmetric, rod-like shape. Due to its size and shape the gynapical chalaza makes a much larger contribution to the overall widening of the chalaza in comparison to the comparably slender gynbasal chalaza. Finally, cells of the gynapical chalaza provide a subepidermal contribution to the outer integument.

## Discussion

Using 3D digital ovules we provide the most comprehensive and detailed cellular description of ovule development in *Arabidopsis thaliana* available to date. To this end we established and extensively optimized the necessary microscopy and segmentation pipeline. The entire procedure from imaging to the final segmented digital ovule takes about 45 minutes per z-stack. For younger ovules the procedure takes even less time. This novel methodology enabled a rapid workflow which allowed the generation of a detailed 3D atlas of an organ of such complex architecture. Importantly, our approach revealed single-cell-level information of deeper tissues of the ovule and allowed simultaneous analysis of multiple fluorescent markers. The 3D digital ovules including their quantitative descriptions can be downloaded from the BioStudies data repository at EMBL-EBI (Sarkans et al., 2018). The experimental strategy is applicable to other organs as well (Tofanelli et al., 2019); Wolny et al., 2020). In an exemplary manner, this work demonstrates the exciting new insights that can be obtained when studying tissue morphogenesis in full 3D.

The unprecedented analytical power inherent in the examination of 3D digital ovules asserted itself at multiple levels. The analysis revealed new insights into the dynamic growth patterns underlying ovule development, identified previously unrecognized cell populations, and provided new information regarding *WUS* expression. 3D digital ovules also enable new insight into gene function by drastically improving the comparative phenotypic analysis between wild-type and mutants. Our data unveiled that *INO* plays an even more important role in the organization of the early ovule than previously appreciated. Indeed, we have found that *ino* mutants not only show a missing outer integument and defective embryo sac development but also exhibit previously unrecognized cellular aberrations in the nucellus and chalaza. It has long been recognized that ontogenesis of the embryo sac correlates with proper integument development although the molecular mechanism remains unclear (Gasser et al., 1998; Grossniklaus and Schneitz, 1998). Our new data allow an alternative interpretation involving hormonal signaling from the chalaza. Cytokinin is known to affect ovule patterning through the control of the expression of the auxin efflux carrier gene *PIN-FORMED 1* (*PIN1*) and a chalaza-localized cytokinin signal is required for early gametophyte development (Bencivenga et al., 2012; Ceccato et al., 2013; Cheng et al., 2013). Thus, the defects in the development of the nucellus and embryo sac of *ino-5* may relate to its cellular misorganization of the subepidermal chalaza and thus altered cytokinin and auxin signaling.

The analysis of 3D digital ovules enabled interesting new insights into central cellular processes that shape the Arabidopsis ovule. Here, we focussed on novel aspects of primordium formation, the initiation and extension of the ii’ layer of the inner integument, and ovule curvature. Regarding primordium formation our data support new notions regarding outgrowth and early pattern formation. Our data indicate that primordium outgrowth is even and does not fluctuate between fast and slow phases of growth. Moreover, we discovered primordium slanting, a hitherto unrecognized aspect of early ovule development. Primordium slanting indicates that gynapical-gynbasal polarity becomes visible morphologically much earlier than previously appreciated. The early appearance of gynapical-gynbasal polarity is in line with molecular data (Sieber et al., 2004).

Our data broadened the repertoire of modes of how a new cell layer can be formed in plants. Cambium-like activity of the innermost layer of the inner integument (endothelium or ii1 layer) results in the formation of the ii1’ layer (Schneitz et al., 1995; Debeaujon et al., 2003). Early morphological analysis in 2D led to the hypothesis that all cells located in the roughly proximal half of the ii1 layer undergo periclinal divisions resulting in the formation of the ii1’ cell layer. This model conformed to the broadly supported notion that periclinal asymmetric cell divisions underlie the formation of new cell layers in plants. For example, in the eight-cell Arabidopsis embryo all cells undergo asymmetric cell divisions and thus contribute to the formation of the epidermis (Jürgens and Mayer, 1994; Yoshida et al., 2014). Similarly, periclinal asymmetric cell divisions in all the radially arranged cortex/endodermis initials are thought to contribute to the formation of the cortex and endodermis cell layers of the main root (Di Laurenzio et al., 1996; Dolan et al., 1993). By contrast, our investigation of 3D digital ovules revealed a distinct mode for the formation of the ii1’ cell layer. The analysis indicated that only a small number of ii1 cells undergo asymmetric cell divisions and produce the first ii1’ cells. Much of subsequent ii1’ layer formation appears to rely on symmetric cell divisions that originate in the daughters of those few ii1’ founder cells. Thus, formation of the ii1’ layer largely depends on lateral extension starting from a few scattered ii1’ pioneer cells, a process that may be regarded as layer invasion. It will be interesting to investigate the molecular control of this new type of cell layer formation in future studies.

We provide novel insight into the internal cellular makeup of the chalaza and how it relates to central aspects of ovule morphogenesis, such as the broadening of the chalaza and kink formation. The morphological comparison of the cellular patterns of the chalaza between wild type and *ino-5* revealed that this region is of complex composition. Our observations indicate that the gynapcial and gynbasal chalaza, two previously unrecognized cell populations of similar size but different shape and cellular composition, contribute to the interior cellular architecture of the proximal central region. In addition, our data suggest that differential growth between the gynapical and gynbasal chalaza results in the bulging of the gynapical chalaza. This process appears to relate to two aspects: the establishment of a group of large cells that in part make a subepidermal contribution to the outer integument and the formation of the “wing-like” flaps in the gynapical chalaza. By contrast, the radially symmetric gynbasal chalaza features a comparably slender shape. Bulging ultimately leads to an arrangement where the nucellus and inner integument appear to sit on top of the gynbasal chalaza.

Finally, we provide new insight into the genetic and cellular processes regulating ovule curvature. Previous studies hypothesized that ovule curvature is a multi-step process with a major involvement by the proximal chalaza and the integuments (Baker et al., 1997; Schneitz et al., 1997). Here, a detailed comparison between wild-type and *ino-5* 3D digital ovules provided evidence for two distinct processes that contribute to ovule curvature: kink formation in the ovule primordium and bending of the developing integuments. Both processes require *INO* function as neither kink formation nor integument bending take place in *ino-5*. The data support the hypothesis that a small number of asymmetric oblique cell divisions in the stage 2-III gynbasal chalaza lead to the characteristic early kink of the ovule primordium. Regarding integument bending it is noteworthy that despite the straight growth of the inner integument of *ino-5* its general cellular parameters appear largely unaffected. In addition, integument bending is prominent in mutants lacking an embryo sac (Schneitz et al., 1997). These observations indicate that the outer integument imposes bending onto the inner integument and nucellus. Future studies will address in more detail the interplay between genetics, cellular behavior, and tissue mechanics that underlies ovule curvature.

## Materials and Methods

### Plant work, plant genetics, and plant transformation

*Arabidopsis thaliana* (L.) Heynh. var. Columbia (Col-0) was used as wild-type strain. Plants were grown as described earlier (Fulton et al., 2009). The Col-0 line carrying the *WUSCHEL* (*WUS*) promoter construct (pWUS::2xVENUS:NLS::tWUS) is equivalent to a previously described reporter line except that vYFP was exchanged for 2xVENUS (Zhao et al., 2017). The *ino-5* mutation (Col-0) was generated using a CRISPR/Cas9 system in which the egg cell-specific promoter pEC1.2 controls Cas9 expression (Wang et al., 2015). The single guide RNA (sgRNA) 5’-ACCATCTATTTGATCTGCCG-3’ binds to the region +34 to +55 of the *INO* coding sequence. The sgRNA was designed according to the guidelines outlined in (Xie et al., 2014). The mutant carries a frameshift mutation at position 51 relative to the *INO* start AUG, which was verified by sequencing. The resulting predicted short INO protein comprises 78 amino acids. The first 17 amino acids correspond to INO while amino acids 18 to 78 represent an aberrant amino acid sequence. Wild-type plants were transformed with different constructs using Agrobacterium strain GV3101/pMP90 (Koncz and Schell, 1986) and the floral dip method (Clough and Bent, 1998). Transgenic T1 plants were selected on Hygromycin (20 μg/ml) plates and transferred to soil for further inspection.

### Recombinant DNA work

For DNA work, standard molecular biology techniques were used. PCR fragments used for cloning were obtained using Q5 high-fidelity DNA polymerase (New England Biolabs, Frankfurt, Germany). All PCR-based constructs were sequenced.

### Clearing and staining of ovules

A detailed protocol was recently published (Tofanelli et al., 2019). Fixing and clearing of dissected ovules in ClearSee was done essentially as described (Kurihara et al., 2015). Tissue was fixed in 4% paraformaldehyde in PBS followed by two washes in PBS before transfer into the ClearSee solution (xylitol (10%, w/v), sodium deoxycholate (15%, w/v), urea (25%, w/v), in H_2_O). Clearing was done at least overnight or for up to two to three days. Staining with SR2200 (Renaissance Chemicals, Selby, UK) was essentially performed as described in (Musielak et al., 2015). Cleared tissue was washed in PBS and then put into a PBS solution containing 0.1% SR2200 and a 1/1000 dilution of TO-PRO-3 iodide (Bink et al., 2001; Van Hooijdonk et al., 1994) (Thermo Fisher Scientific) for 20 minutes. Tissue was washed in PBS for one minutes, transferred again to ClearSee for 20 minutes before mounting in Vectashield antifade agent (Florijn et al., 1995) (Vectashield Laboratories, Burlingame, CA, USA).

### Microscopy and image acquisition

Confocal laser scanning microscopy of ovules stained with SR2200 and TO-PRO-3 iodide was performed on an upright Leica TCS SP8 X WLL2 HyVolution 2 (Leica Microsystems) equipped with GaAsP (HyD) detectors and a 63x glycerol objective (HC PL APO CS2 63x/1.30 GLYC, CORR CS2). Scan speed was at 400 Hz, line average between 2 and 4, and the digital zoom between 1 and 2. Laser power or gain was adjusted for z compensation to obtain an optimal z-stack. SR2200 fluorescence was excited with a 405 nm diode laser (50 mW) with a laser power ranging from 0.1 to 1.5 percent intensity and detected at 420 to 500 nm with the gain of the HyD detector set to 20. TO-PRO-3 iodide fluorescence excitation was done at 642 nm with the white-light laser, with a laser power ranging from 2 to 3.5 percent and detected at 655 to 720 nm, with the gain of the HyD detector set to 200. For high quality z-stacks, 12-bit images were captured at a slice interval of 0.24 μm with optimized system resolution of 0.063 μm × 0.063 μm × 0.240 μm as final pixel size according to the Nyquist criterion. Some of the z stacks were captured with 2x downsampled voxel size of 0.125 μm × 0.125 μm × 0.24 μm where we took advantage of the PlantSeg-trained model to generate equal standard cell segmentation as with images with fine voxels. This was possible because the PlantSeg model “generic_confocal_3d_unet” was trained on downsampled original images and ground truths. The model now requires raw images whose voxels are scaled to the trained dataset so that it generates best cell boundary predictions. Overall, raw images captured with 2x downsampled voxels were helpful in that it simplified the rescaling step in PlantSeg and allowed us to generate raw images in less time without compromising segmentation quality. Scan speed was set to 400 Hz, the pinhole was set to 0.6 Airy units, line average was between 2 and 4, and the digital zoom was set between 1 and 2, as required. Laser power or gain was adjusted for z compensation to obtain an optimal z-stack. Images were adjusted for color and contrast using Adobe Photoshop CC (Adobe, San Jose, CA, USA), GIMP (https://www.gimp.org/) or MorphographX (Barbier de Reuille et al., 2015) software (https://www.mpiz.mpg.de/MorphoGraphX). Image acquisition parameters for pWUS::2xVenus reporter line were the following: SR2200; 405 diode laser 0.10%, HyD 420–480 nm, detector gain 10. 2xVenus; 514 nm Argon laser 2%, HyD 525-550 nm, detector gain 100. TO-PRO-3; 642 nm White Laser 2%, HyD 660–720 nm, detector gain 100. In each case sequential scanning was performed to avoid crosstalk between the spectra.

### Segmentation

#### 3D cell segmentation

3D cells were segmented using PlantSeg (Wolny et al. 2020). Cell segmentation using boundary predictions in PlantSeg allowed us to process microscopic image z-stacks captured with coarse voxels, to minimise the segmentation errors, and to reduce the imaging time and hence laser exposure and phototoxicity to samples. The PlantSeg pipeline takes as input a batch of raw images in tiff format depicting the cell walls stained by SR2200 and outputs cell boundary predictions by a U-Net-based convolutional neural network (CNN) (Ronneberger et al., 2015) together with 3D cell segmentation given by partitioning of the boundary predictions. The PlantSeg pipeline is subdivided in four sequential steps. The first step consists of pre-processing the raw files. In particular, we found rescaling the raw images to match the U-Net training data crucial to achieve the best performances. This can be done semi-automatically (see below) by using the guided rescaling tool in the data pre-processing section of the PlantSeg gui. The output of the pre-processing step is saved as hierarchical data format (HDF5). The second step consists of predicting the cell boundaries. PlantSeg has built-in several pre-trained CNNs that can be chosen for different types of raw input data. For boundary prediction the “generic_confocal_3D_unet” CNN was employed, which was trained on the Arabidopsis ovules dataset (https://osf.io/w38uf). The third step is where the actual segmentation is obtained. PlantSeg’s segmentation is based on graph partitioning and one can choose between several partitioning strategies, such as Multicut (Speth et al., 2011) or GASP average (Bailoni et al., 2019). The last step deals with image post processing. This step is necessary to convert the images to the original voxel resolution and to convert the results from HDF5 back to tiff file format. A thorough experimental evaluation of the different parameters revealed that optimal results could be obtained by using voxel size 0.15 × 0.15 × 0.235 (μm, xyz) for the guided rescaling, “generic_confocal_3D_unet” CNN for cell boundary prediction, and cell segmentation performed by GASP average and watershed in 3D (Bailoni et al. 2019) with default parameters. The PlantSeg’s YAML configuration file with all the parameter settings for the pipeline is found in the Supplementary File 1. These settings allowed for near-perfect 3D cell segmentation as only a small number of errors, such as over-segmented cells, had to be corrected by visual inspection of the segmented stack in MGX. In critical cases this included cross-checking the TO-PRO-3 channel which included the stained nuclei. Thus, we now routinely image ovules using these settings. Image acquisition of mature ovules takes about 15 minutes for both channels (SR2200/cell contours; TO-PRO-3/nuclei), running the PlantSeg pipeline requires about 25 minutes on our computer hardware (1x Nvidia Quadro P5000 GPU), and manual correction of segmentation errors takes less than five minutes.

#### *Quantifying* pWUS *nuclei in the ovule*

The nuclei labelled by pWUS signal (blobs) were counted using the `Local Maxima` process in MGX. The raw z-stacks were processed with the process `Stack/Filter/Brighten Darken` with a value ranging from 2 to 4 depending on image quality. Further Gaussian blur was applied twice with a low sigma of 0.2 × 0.2 × 0.2 μm^3^ using `Stack/Filter/Gaussian Blur Stack`. The processed stack was used to generate local signal maxima of radius 1.5 μm in xyz and with a typical threshold of 8000 using process `Stack/Segmentation/Local Maxima`. The blobs were visualised by creating 3D blob meshes using the process `Mesh/Creation/Mesh From Local Maxima`. Blob size can be adjusted as required. We used a blob radius of size 1.2 μm that fits inside the nuclei. The blobs were also linked to their parent 3D cell meshes using the process `Mesh/Nucleus/Label Nuclei`. The blob meshes were further linked to their parent 3D cell meshes using the MGX process Mesh/Nucleus/Label Nuclei. This requires the 3D cell meshes to be loaded into the `mesh 1` workspace and the blob meshes into the `mesh 2` workspace of MGX. This process sets the cell identities of the 3D cell meshes as parents to the blob meshes. Blob numbers were determined in exported .csv files that contained the Blob IDs associated with their parent cell IDs.

### Generation of 3D Cell meshes and classification of cell types

3D cell meshes were generated in MGX using the segmented image stacks using the process `Mesh/Creation/Marching Cube 3D` with a cube size of 1. All the cell annotation was performed on the 3D cell meshes in MGX. Tissue surface mesh is required for the method of semi-automatic cell type labelling. Tissue surface mesh is generated from the segmented stack. The segmented stack was first gaussian blurred using the process `Stack/Filter/Gaussian Blur Stack` with a radius of 0.3 in xyz. The smooth stack was used to generate tissue surface mesh using `Mesh/Creation/Marching Cube Surface` with a cube size of 1 and threshold 1. The generated surface mesh was then smoothed several times using the process `Mesh/Structure/Smooth mesh` with 10 passes. For mature ovules cell type annotation, we used the MGX process `Mesh/Cell Atlas 3D/Ovule/Detect Cell Layer` (a modified 3DCellAtlas Meristem tool (Montenegro-Johnson et al., 2019)) with the 3D cell meshes in the `mesh 1` workspace and the tissue surface mesh in the `mesh 2` workspace using a cone angle parameter of 1.2. This process correctly classified about 60 percent of cells based on the layer they belong to. We manually corrected mis-annotations and labelled the rest of the cells by using the mesh tools in MGX. We used the `Select Connected Area` tool to select individual cells of different layers in 3D and proofread the cell type labelling with `Mesh/Cell Types/Classification/Set Cell Type`. Each cell layer of the integuments was consecutively shaved off and proofreading was performed using 3D surface view. We further used the processes `Mesh/Cell Types/Classification` to save all labels, load labels and select cell types as required. The saved cell types.csv file was reloaded onto the original 3D cell mesh and final proofreading was performed in the section view. For primordia we manually annotated the cells using the tools in `Mesh/Cell Types/Classification`.

### Exporting attributes from MorphographX for further quantitative analysis

All quantitative cellular features were exported as attributes from MGX. The attributes included cell IDs (segmentation label of individual cells), cell type IDs (tissue annotation), and cell volume. The attributes from individual ovules were exported as .csv files and merged to create long-format Excel-sheets listing all the scored attributes of all the cells from the analyzed ovules. These files are included in the downloadable datasets.

### Computer requirements

A minimal setup requires an Intel Core i9 CPU with at least 64 GB of RAM. To take full advantage of the PlantSeg pipeline requires at a minimum an 8 GB NVIDIA GPU/CUDA graphics card (for example a Geforce RTX 2080). For routine work we use a computer with a 16 GB NVIDIA QUADRO P5000 card. We use Linux Mint as the operating system (www.linuxmint.com).

### Growth rate and relative tissue growth

Stage-specific growth rates were estimated by taking the ratio between a given mean parameter of two consecutive stages (x(t+Δt)/x(t)). Relative growth of an ovule tissue was estimated by taking the ratio of the respective mean parameter of two consecutive stages divided by the ovule growth rate (y(t+Δt)/y(t)/(x(t+Δt)/x(t)).

### Primordia length and slanting

Length along the surface of the gynapical and gynbasal sides of primordia was quantified for individual primordia. We extracted a file of epidermal cells at the mid-section of two halves of the ovule and placed a Bezier grid of size 3 × 3 (xy) on the distal tip of the primordia surface. We then used the MGX process “Mesh/Heatmap/Measure 3D/Distance to Bezier” to quantify the length. The measured values are the distance from the Bezier grid to individual cell centroids through the file of connected cells. Primordia height was quantified by averaging the two values. Slanting was quantified by obtaining the difference of the maximal values at the gynapical and gynbasal sides, respectively (Supplementary Fig. 1).

### Statistical analysis

Statistical analysis was performed using a combination of R (R Core Team, 2019) with RStudio (RStudio Team, 2019), the Anaconda distribution (Anaconda Software Distribution; https://anaconda.com) of the Python SciPy software stack (Oliphant, 2007), and PRISM8 software (GraphPad Software, San Diego, USA).

### Datasets

The entire 3D digital ovule datasets can be downloaded from the BioStudies data repository (https://www.ebi.ac.uk/biostudies/). Each dataset contains raw cell boundaries, cell boundaries, predictions from PlantSeg, nuclei images, segmented cells as well as the annotated 3D cell meshes, and the associated attribute files in csv format. The 3D mesh files can be opened in MGX. Accession S-BSST475: the wild-type high-quality dataset and the additional dataset with more segmentation errors. Accession S-BSST498: pWUS::2xVenus:NLS. Accession S-BSST497: *ino-5*. Accession S-BSST513: Long-format Excel-sheets listing all the scored attributes of all the cells from the analyzed ovules.

## Acknowledgements

We thank members of the Schneitz lab for helpful comments. We also thank Lynette Fulton, Thomas Greb, and Alexis Maizel for input and comments on the manuscript. We further thank Miltos Tsiantis for comments and support. We appreciate the gift of the pWUS line by Christian Wenzel and Jan Lohmann. We further acknowledge support by the Center for Advanced Light Microscopy (CALM) of the TUM School of Life Sciences. This work was funded by the German Research Council (DFG) through grants FOR2581 (TP3) to AK and FH, (TP8) to RS, and (TP7) to KS.

## Authors’ contributions

AV, RT, RS and KS designed the study. AV, RT, SS, LC and AW performed the experiments. RT, AV, SS, LC, AW, AK, FH, RS and KS interpreted the results. AK, FH, RS and KS secured funding. KS wrote the paper. All authors read and approved the final manuscript.

## Supplementary Materials

**Supplementary Table 1.**
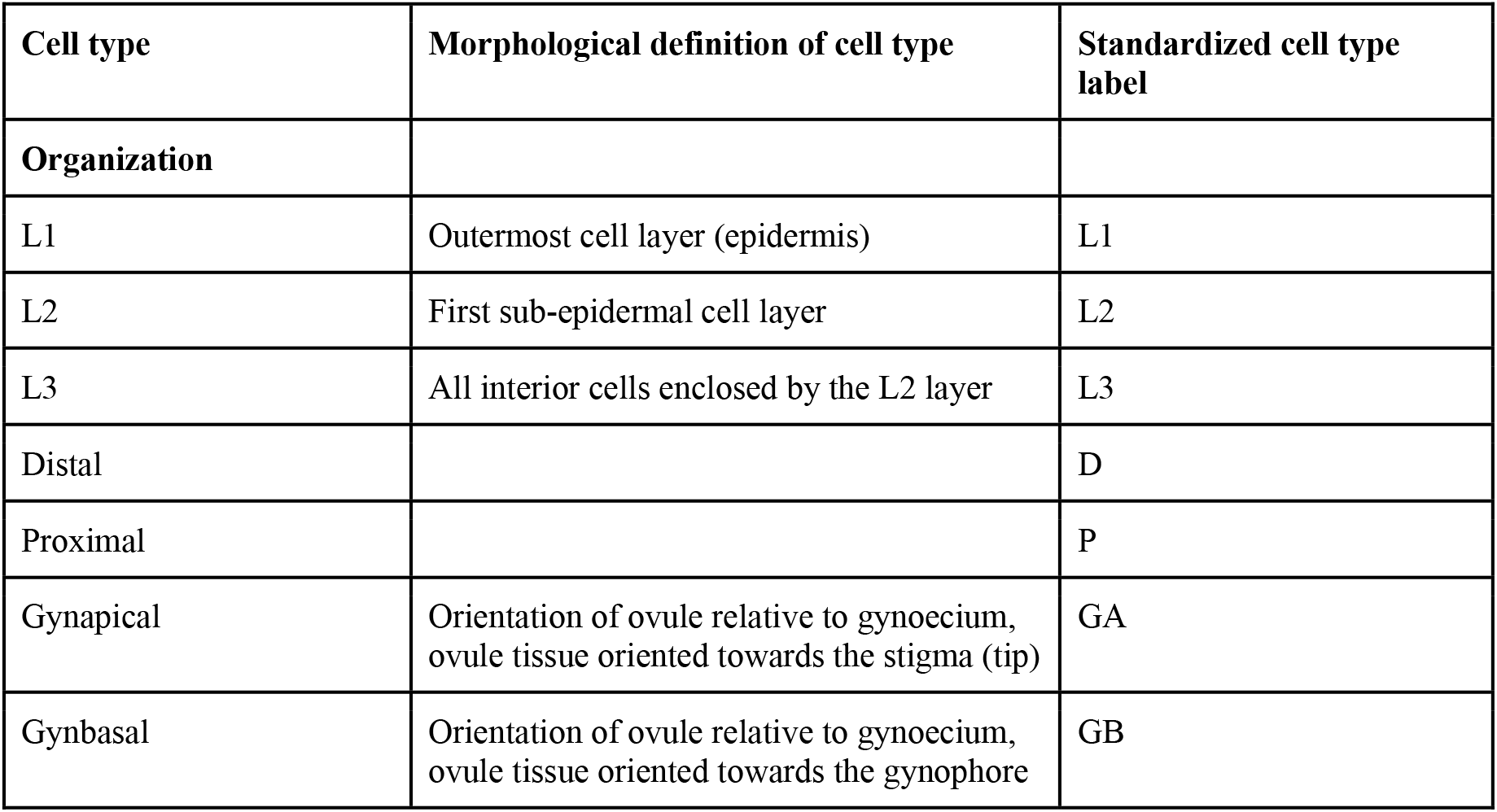
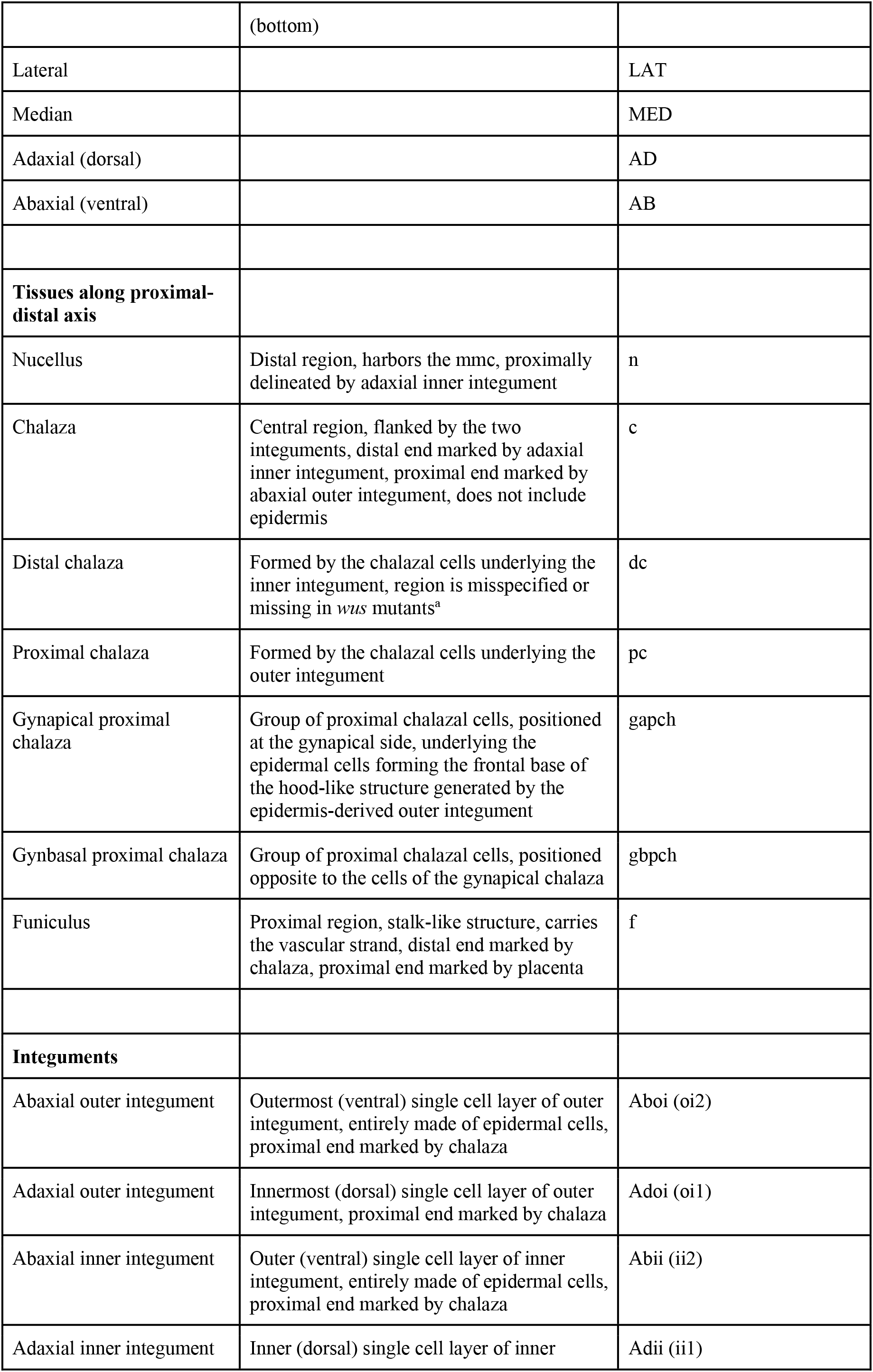
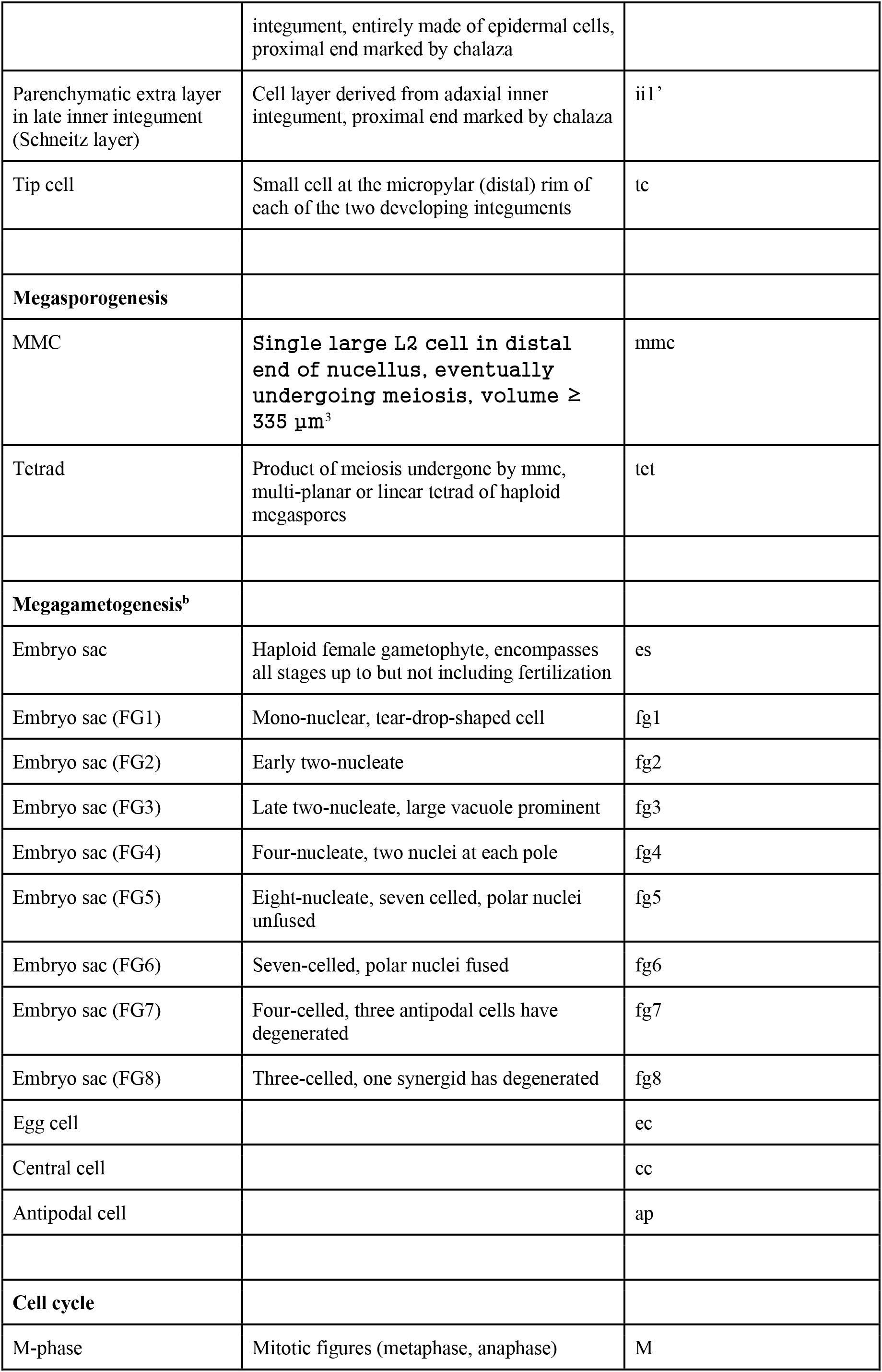
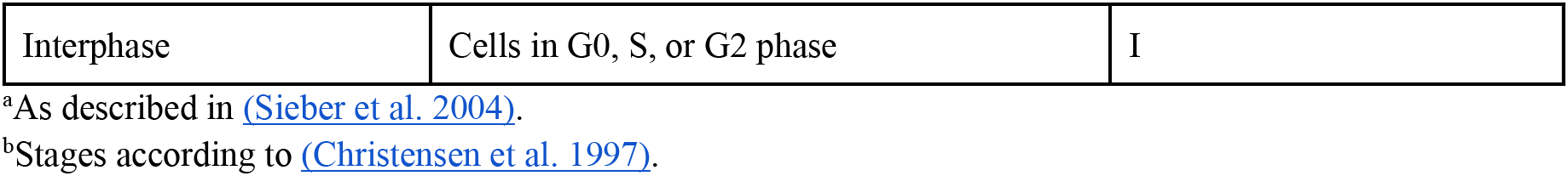
Standardized cell type labels.

| Cell type | Morphological definition of cell type | Standardized cell type label |
| --- | --- | --- |
| **Organization** |  |  |
| L1 | Outermost cell layer (epidermis) | L1 |
| L2 | First sub-epidermal cell layer | L2 |
| L3 | All interior cells enclosed by the L2 layer | L3 |
| Distal |  | D |
| Proximal |  | P |
| Gynapical | Orientation of ovule relative to gynoecium, ovule tissue oriented towards the stigma (tip) | GA |
| Gynbasal | Orientation of ovule relative to gynoecium, ovule tissue oriented towards the gynophore | GB |
|  | (bottom) |  |
| Lateral |  | LAT |
| Median |  | MED |
| Adaxial (dorsal) |  | AD |
| Abaxial (ventral) |  | AB |
| **Tissues along proximal-distal axis** |  |  |
| Nucellus | Distal region, harbors the mmc, proximally delineated by adaxial inner integument | n |
| Chalaza | Central region, flanked by the two integuments, distal end marked by adaxial inner integument, proximal end marked by abaxial outer integument, does not include epidermis | c |
| Distal chalaza | Formed by the chalazal cells underlying the inner integument, region is misspecified or missing in *wus* mutants<sup>a</sup> | dc |
| Proximal chalaza | Formed by the chalazal cells underlying the outer integument | pc |
| Gynapical proximal chalaza | Group of proximal chalazal cells, positioned at the gynapical side, underlying the epidermal cells forming the frontal base of the hood-like structure generated by the epidermis-derived outer integument | gapch |
| Gynbasal proximal chalaza | Group of proximal chalazal cells, positioned opposite to the cells of the gynapical chalaza | gbpch |
| Funiculus | Proximal region, stalk-like structure, carries the vascular strand, distal end marked by chalaza, proximal end marked by placenta | f |
| **Integuments** |  |  |
| Abaxial outer integument | Outermost (ventral) single cell layer of outer integument, entirely made of epidermal cells, proximal end marked by chalaza | Aboi (oi2) |
| Adaxial outer integument | Innermost (dorsal) single cell layer of outer integument, proximal end marked by chalaza | Adoi (oi1) |
| Abaxial inner integument | Outer (ventral) single cell layer of inner integument, entirely made of epidermal cells, proximal end marked by chalaza | Abii (ii2) |
| Adaxial inner integument | Inner (dorsal) single cell layer of inner | Adii (ii1) |
|  | integument, entirely made of epidermal cells, proximal end marked by chalaza |  |
| Parenchymatic extra layer in late inner integument (Schneitz layer) | Cell layer derived from adaxial inner integument, proximal end marked by chalaza | ii1' |
| Tip cell | Small cell at the micropylar (distal) rim of each of the two developing integuments | tc |
| **Megasporogenesis** |  |  |
| MMC | Single large L2 cell in distal end of nucellus, eventually undergoing meiosis, volume $\geq 335 \mu\text{m}^3$ | mmc |
| Tetrad | Product of meiosis undergone by mmc, multi-planar or linear tetrad of haploid megaspores | tet |
| **Megagametogenesis<sup>b</sup>** |  |  |
| Embryo sac | Haploid female gametophyte, encompasses all stages up to but not including fertilization | es |
| Embryo sac (FG1) | Mono-nuclear, tear-drop-shaped cell | fg1 |
| Embryo sac (FG2) | Early two-nucleate | fg2 |
| Embryo sac (FG3) | Late two-nucleate, large vacuole prominent | fg3 |
| Embryo sac (FG4) | Four-nucleate, two nuclei at each pole | fg4 |
| Embryo sac (FG5) | Eight-nucleate, seven celled, polar nuclei unfused | fg5 |
| Embryo sac (FG6) | Seven-celled, polar nuclei fused | fg6 |
| Embryo sac (FG7) | Four-celled, three antipodal cells have degenerated | fg7 |
| Embryo sac (FG8) | Three-celled, one synergid has degenerated | fg8 |
| Egg cell |  | ec |
| Central cell |  | cc |
| Antipodal cell |  | ap |
| **Cell cycle** |  |  |
| M-phase | Mitotic figures (metaphase, anaphase) | M |
| Interphase | Cells in G0, S, or G2 phase | I |

<sup>a</sup>As described in (Sieber et al. 2004).
<sup>b</sup>Stages according to (Christensen et al. 1997).

**Supplementary Table 2.** Cell numbers and total volumes of *ino-5* ovules staged according to the wild-type cohort with removed outer integument.

| Stage<sup>a,b</sup> | N Cells wild type (-oi) | Volume (x10<sup>4</sup> μm<sup>3</sup>) wild type (-oi) | N Cells *ino-5* | Volume (x10<sup>4</sup> μm<sup>3</sup>) *ino-5* |
| --- | --- | --- | --- | --- |
| 1-II | 74.0 ± 17.1 | 1.0 ± 0.2 | 87 ± 23.1 | 1.1 ± 0.3 |
| 2-I | 176.9 ± 31.5 | 2.5 ± 0.4 | 166.2 ± 19.5 | 2.1 ± 0.1 |
| 2-II | 220.6 ± 24.9 | 2.7 ± 0.6 | 241.1 ± 9.1 | 3.0 ± 0.2 |
| 2-III | 270.3 ± 32.4 | 3.2 ± 0.6 | 305.5 ± 34.6 | 3.7 ± 0.3 |
| 2-IV | 380.5 ± 26.5 | 4.5 ± 0.4 | 382.3 ± 45.7 | 4.1 ± 0.5 |
| 2-V | 524.7 ± 66.3 | 6.2 ± 1.0 | 536.7 ± 60.8 | 6.4 ± 0.9 |
| 3-I | 712.2 ± 61.2 | 9.2 ± 1.1 | 716.9 ± 58.8 | 9.4 ± 0.9 |
| 3-II | 888.4 ± 52.9 | 13.8 ± 1.3 | 858.0 ± 28.7 | 12.7 ± 0.9 |
| 3-III | 951.2 ± 90.1 | 14.9 ± 2.1 | 956.7 ± 28.9 | 15.5 ± 0.6 |
| 3-IV | 1037 ± 103.7 | 18.1 ± 2.2 | 1102 ± 49.5 | 19.6 ± 2.1 |
| 3-V | 1140 ± 96.8 | 20.8 ± 2.3 | 1241 ± 32.2 | 23.5 ± 17.7 |
| 3-VI | 1345 ± 131.9 | 26.6 ± 3.7 | 1383 ± 61.1 | 25.7 ± 16.8 |

<sup>a</sup>Number of 3D digital ovules scored: 10 (stages 2-II- 3-II, 3-IV, 3-VI), 11 (stages 2-I, 3-III, 3-V), 13 (stage 2-II), 14 (stage 1-I), 28 (stage 1-II).
Values represent mean ± SD.
<sup>b</sup>Number of 3D digital ovules scored for *ino-5* dataset: 3 (stages 2-IV), 6 (stages 2-III), 7 (stages 2-II, 3-III, 3-VI), 9 (stages 3-II), 10 (stages 2-I, 3-V), 12 (stages 1-II), 14 (stages 3-I, 3-IV), 20 (stages 2-V).

**Supplementary Fig 1.**
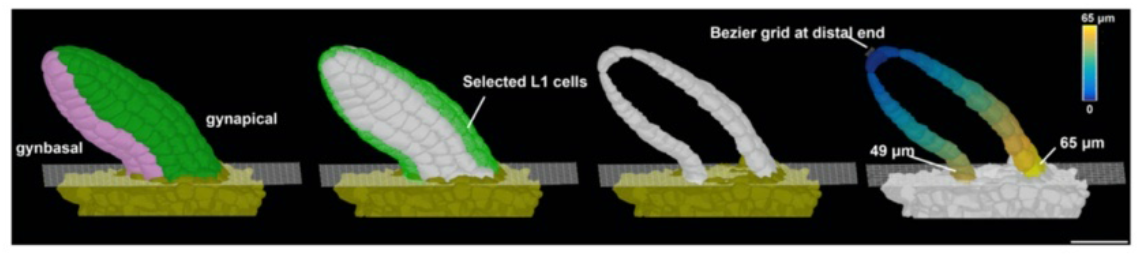
Primordium length and slant measurement. 3D surface view of primordia displaying the method for length measurement. The heatmap on the extracted cells depicts the quantified distance value between individual measured cells to the distal tip of primordia. Scale bars: 20 μm.

**Supplementary File 1. Yaml file with PlantSeg parameters.**

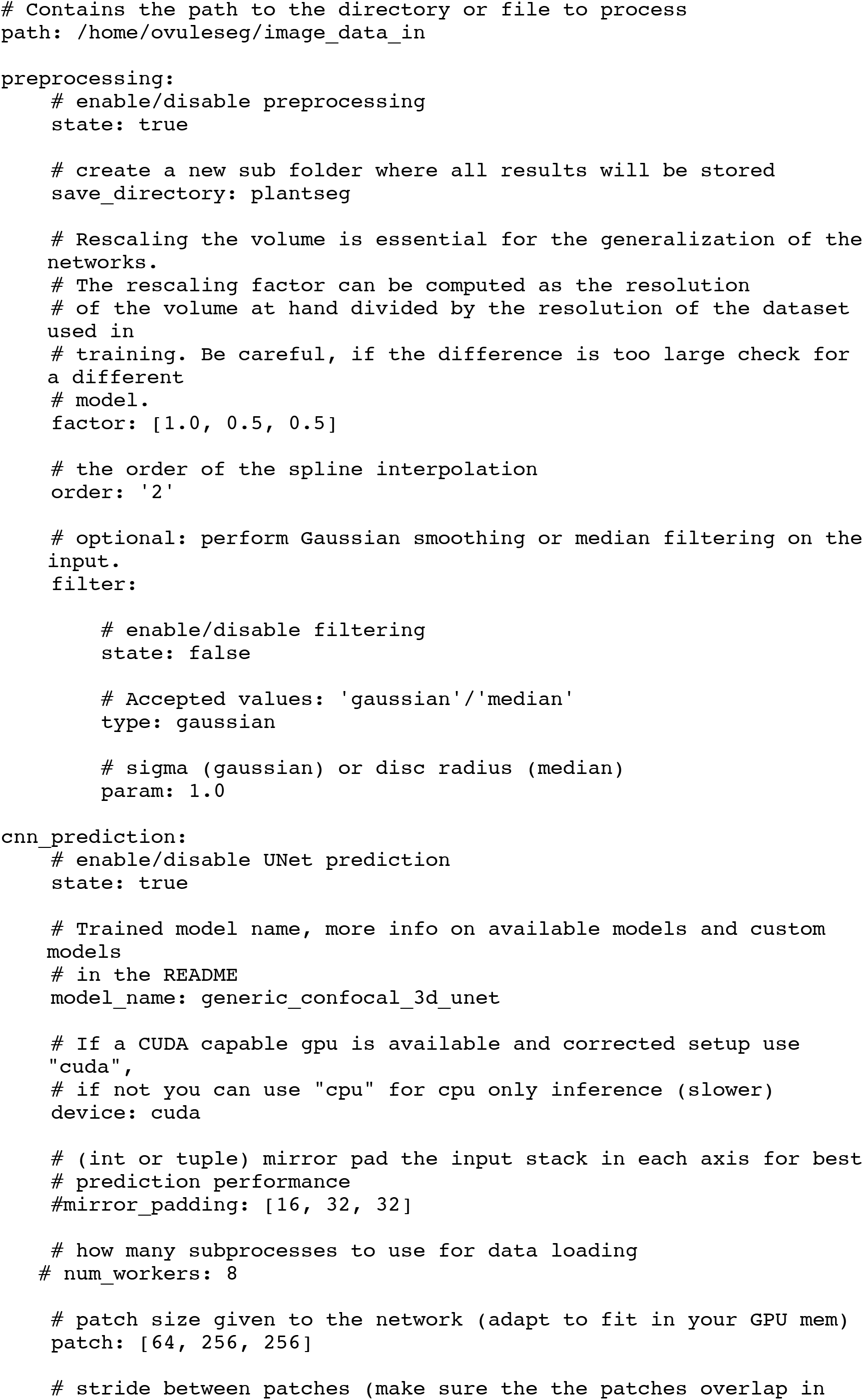

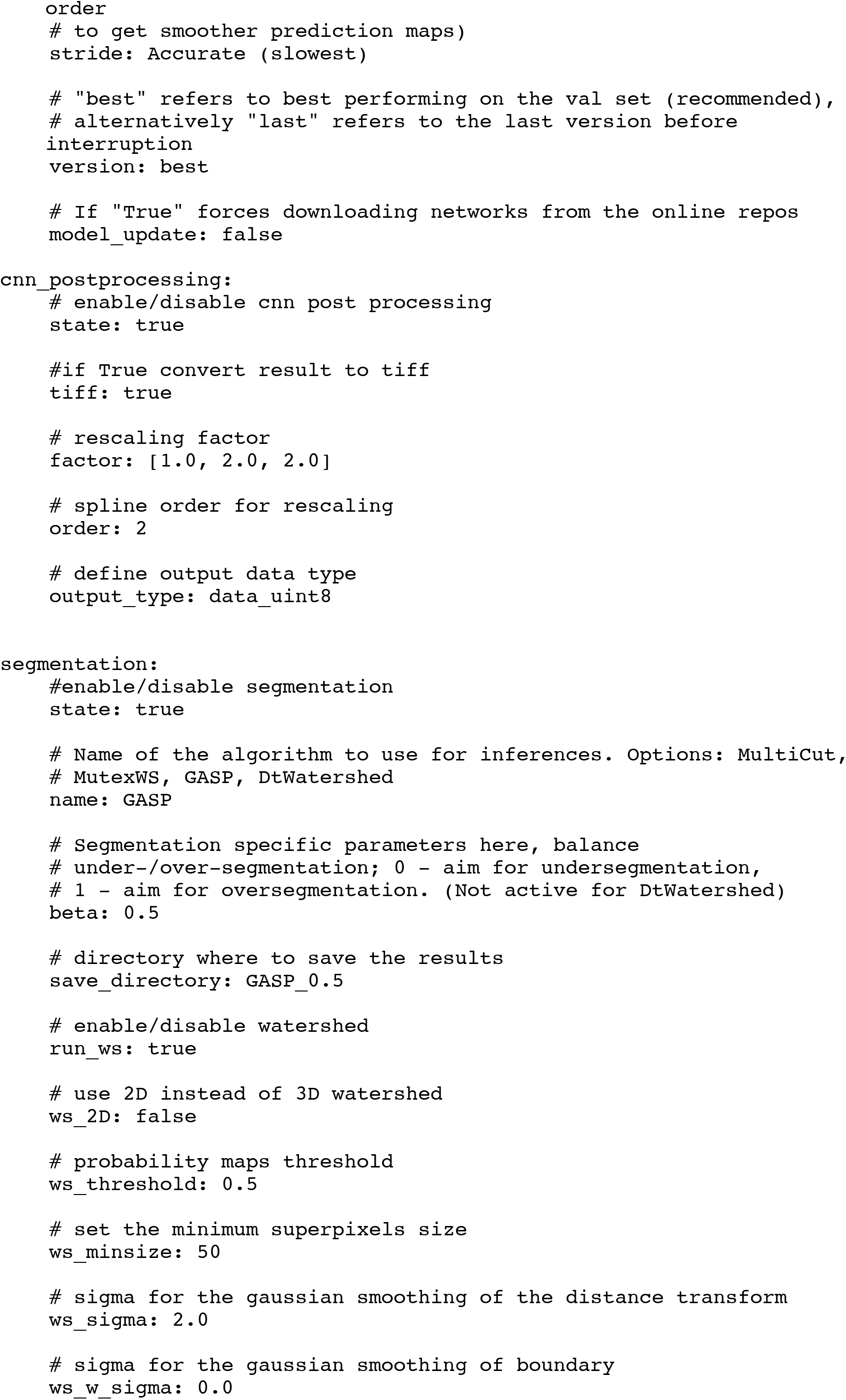

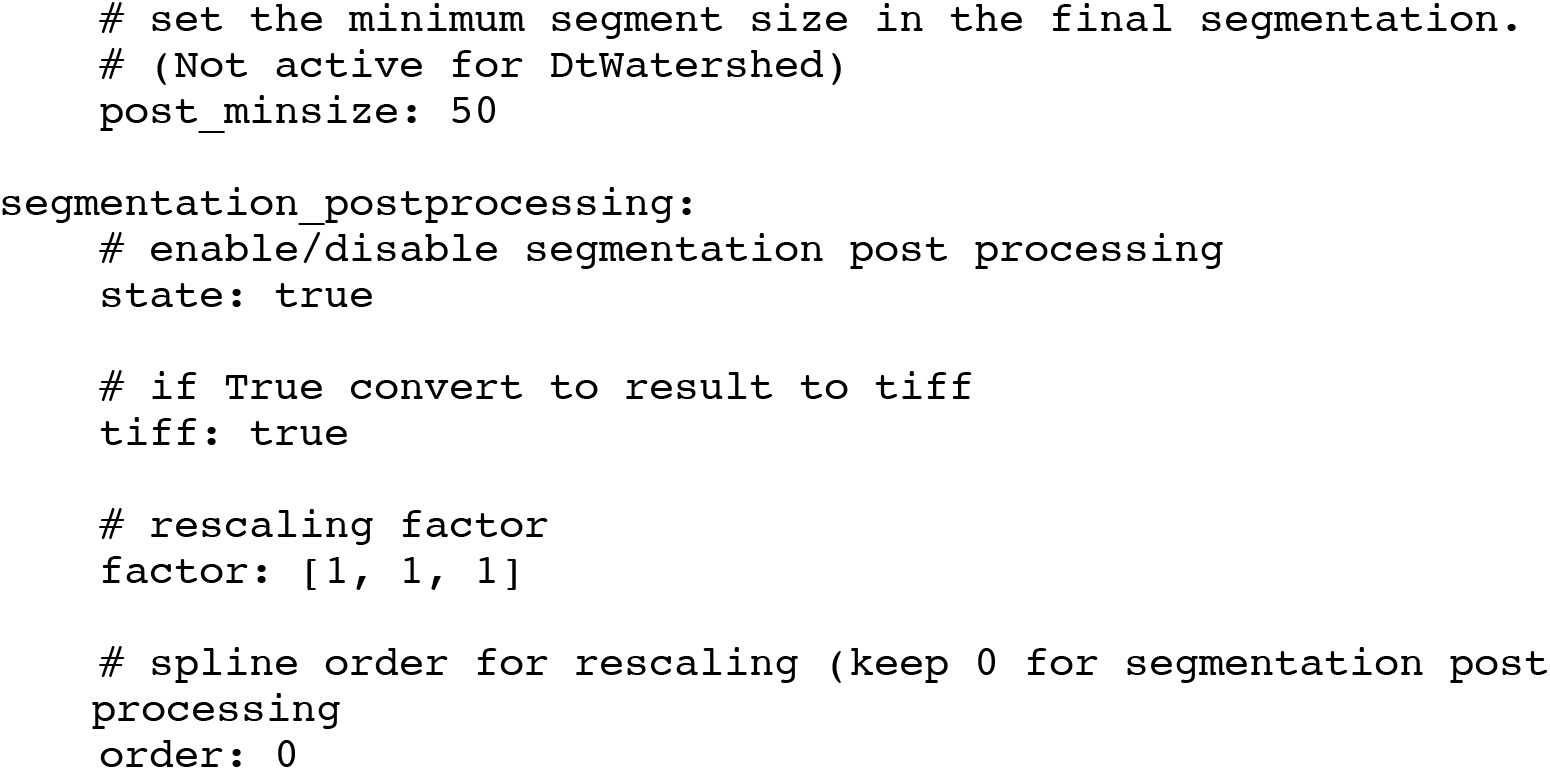

## References

Anderson, C., Hill, B., Lu, H.-C., Moverley, A., Yang, Y., Oliveira, N.M.M., Baldock, R.A., Stern, C.D., 2019. A 3D molecular atlas of the chick embryonic heart. Dev. Biol. 456, 40–46. doi:10.1016/j.ydbio.2019.07.003

Asadulina, A., Panzera, A., Verasztó, C., Liebig, C., Jékely, G., 2012. Whole-body gene expression pattern registration in *Platynereis* larvae. Evodevo 3, 27. doi:10.1186/2041-9139-3-27

Bailoni, A., Pape, C., Wolf, S., Beier, T., Kreshuk, A., Hamprecht, F.A., 2019. A generalized framework for agglomerative clustering of signed graphs applied to instance segmentation. https://arxiv.org/abs/1906.11713.

Baker, S.C., Robinson-Beers, K., Villanueva, J.M., Gaiser, J.C., Gasser, C.S., 1997. Interactions among genes regulating ovule development in. Genetics 145, 1109–1124.

Balasubramanian, S., Schneitz, K., 2000. NOZZLE regulates proximal-distal pattern formation, cell proliferation and early sporogenesis during ovule development in Arabidopsis thaliana. Development 127, 4227–4238.

Balasubramanian, S., Schneitz, K., 2002. *NOZZLE* links proximal-distal and adaxial-abaxial pattern formation during ovule development in *Arabidopsis thaliana*. Development 129, 4291–4300.

Barbier de Reuille, P., Routier-Kierzkowska, A.-L., Kierzkowski, D., Bassel, G.W., Schüpbach, T., Tauriello, G., Bajpai, N., Strauss, S., Weber, A., Kiss, A., Burian, A., Hofhuis, H., Sapala, A., Lipowczan, M., Heimlicher, M.B., Robinson, S., Bayer, E.M., Basler, K., Koumoutsakos, P., Roeder, A.H.K., Smith, R.S., 2015. MorphoGraphX: A platform for quantifying morphogenesis in 4D. elife 4, 05864. doi:10.7554/eLife.05864

Bassel, G.W., Stamm, P., Mosca, G., Barbier de Reuille, P., Gibbs, D.J., Winter, R., Janka, A., Holdsworth, M.J., Smith, R.S., 2014. Mechanical constraints imposed by 3D cellular geometry and arrangement modulate growth patterns in the Arabidopsis embryo. Proc Natl Acad Sci USA 111, 8685–8690. doi:10.1073/pnas.1404616111

Beeckman, T., De Rycke, R., Viane, R., Inzé, D., 2000. Histological study of seed coat development in *Arabidopsis thaliana*. J. Plant Res. 113, 139–148. doi:10.1007/PL00013924

Bencivenga, S., Simonini, S., Benková, E., Colombo, L., 2012. The transcription factors BEL1 and SPL are required for cytokinin and auxin signaling during ovule development in Arabidopsis. Plant Cell 24, 2886–2897. doi:10.1105/tpc.112.100164

Bink, K., Walch, A., Feuchtinger, A., Eisenmann, H., Hutzler, P., Höfler, H., Werner, M., 2001. TO-PRO-3 is an optimal fluorescent dye for nuclear conterstaining in dual-colour FISH on paraffin sections. Histochem Cell Biol 115, 292–299.

Boutros, M., Heigwer, F., Laufer, C., 2015. Microscopy-based high-content screening. Cell 163, 1314–1325. doi:10.1016/j.cell.2015.11.007

Ceccato, L., Masiero, S., Sinha Roy, D., Bencivenga, S., Roig-Villanova, I., Ditengou, F.A., Palme, K., Simon, R., Colombo, L., 2013. Maternal control of PIN1 is required for female gametophyte development in Arabidopsis. PLoS ONE 8, e66148. doi:10.1371/journal.pone.0066148

Chaudhary, A., Gao, J., Schneitz, K., 2018. The genetic control of ovule development, in: Reference Module in Life Sciences. Elsevier. doi:10.1016/B978-0-12-809633-8.20737-1

Cheng, C.-Y., Mathews, D.E., Schaller, G.E., Kieber, J.J., 2013. Cytokinin-dependent specification of the functional megaspore in the Arabidopsis female gametophyte. Plant J. 73, 929–940. doi:10.1111/tpj.12084

Christensen, C.A., King, E.J., Jordan, J.R., Drews, G.N., 1997. Megagametogenesis in Arabidopsis wild type and the Gf mutant. Sex. Plant Reprod. 10, 49–64. doi:10.1007/s004970050067

Clough, S.J., Bent, A.F., 1998. Floral dip: a simplified method for *Agrobacterium*-mediated transformation of *Arabidopsis thaliana*. Plant J. 16, 735–743. doi:10.1046/j.1365-313x.1998.00343.x

Coen, E., Kennaway, R., Whitewoods, C., 2017. On genes and form. Development 144, 4203–4213. doi:10.1242/dev.151910

Coen, E., Rebocho, A.B., 2016. Resolving conflicts: modeling genetic control of plant morphogenesis. Dev. Cell 38, 579–583. doi:10.1016/j.devcel.2016.09.006

Debeaujon, I., Nesi, N., Perez, P., Devic, M., Grandjean, O., Caboche, M., Lepiniec, L., 2003. Proanthocyanidin-accumulating cells in Arabidopsis testa: regulation of differentiation and role in seed development. Plant Cell 15, 2514–2531. doi:10.1105/tpc.014043

Di Laurenzio, L., Wysocka-Diller, J., Malamy, J.E., Pysh, L., Helariutta, Y., Freshour, G., Hahn, M.G., Feldmann, K.A., Benfey, P.N., 1996. The SCARECROW gene regulates an asymmetric cell division that is essential for generating the radial organization of the Arabidopsis root. Cell 86, 423–433. doi:10.1016/s0092-8674(00)80115-4

Dolan, L., Janmaat, K., Willemsen, V., Linstead, P., Poethig, S., Roberts, K., Scheres, B., 1993. Cellular organisation of the *Arabidopsis thaliana* root. Development 119, 71–84.

Dreyer, D., Vitt, H., Dippel, S., Goetz, B., El Jundi, B., Kollmann, M., Huetteroth, W., Schachtner, J., 2010. 3D standard brain of the red flour beetle *Tribolium castaneum*: A tool to study metamorphic development and adult plasticity. Front. Syst. Neurosci. 4, 3. doi:10.3389/neuro.06.003.2010

Endress, P.K., 2011. Angiosperm ovules: diversity, development, evolution. Ann. Bot. 107, 1465–1489. doi:10.1093/aob/mcr120

Florijn, R.J., Slats, J., Tanke, H.J., Raap, A.K., 1995. Analysis of antifading reagents for fluorescence microscopy. Cytometry 19, 177–182. doi:10.1002/cyto.990190213

Fulton, L., Batoux, M., Vaddepalli, P., Yadav, R.K., Busch, W., Andersen, S.U., Jeong, S., Lohmann, J.U., Schneitz, K., 2009. DETORQUEO, QUIRKY, and ZERZAUST represent novel components involved in organ development mediated by the receptor-like kinase STRUBBELIG in Arabidopsis thaliana. PLoS Genet. 5, e1000355. doi:10.1371/journal.pgen.1000355

Gaillochet, C., Daum, G., Lohmann, J.U., 2015. O cell, where art thou? The mechanisms of shoot meristem patterning. Curr. Opin. Plant Biol. 23, 91–97. doi:10.1016/j.pbi.2014.11.002

Gasser, C.S., Broadhvest, J., Hauser, B.A., 1998. Genetic analysis of ovule development. Annu. Rev. Plant Physiol. Plant Mol. Biol. 49, 1–24. doi:10.1146/annurev.arplant.49.1.1

Gasser, C.S., Skinner, D.J., 2019. Development and evolution of the unique ovules of flowering plants. Curr. Top. Dev. Biol. 131, 373–399. doi:10.1016/bs.ctdb.2018.10.007

Gibson, W.T., Veldhuis, J.H., Rubinstein, B., Cartwright, H.N., Perrimon, N., Brodland, G.W., Nagpal, R., Gibson, M.C., 2011. Control of the mitotic cleavage plane by local epithelial topology. Cell 144, 427–438. doi:10.1016/j.cell.2010.12.035

Gross-Hardt, R., Lenhard, M., Laux, T., 2002. *WUSCHEL* signaling functions in interregional communication during *Arabidopsis* ovule development. Genes Dev. 16, 1129–1138. doi:10.1101/gad.225202

Grossniklaus, U., Schneitz, K., 1998. The molecular and genetic basis of ovule and megagametophyte development. Seminars in Cell and Developmental Biology 9, 227–238.

Guignard, L., Fiúza, U.M., Leggio, B., Laussu, J., Faure, E., Michelin, G., Biasuz, K., Hufnagel, L., Malandain, G., Godin, C., Lemaire, P., 2020. Contact area-dependent cell communication and the morphological invariance of ascidian embryogenesis. Science 369. doi:10.1126/science.aar5663

Hamant, O., Heisler, M.G., Jönsson, H., Krupinski, P., Uyttewaal, M., Bokov, P., Corson, F., Sahlin, P., Boudaoud, A., Meyerowitz, E.M., Couder, Y., Traas, J., 2008. Developmental patterning by mechanical signals in Arabidopsis. Science 322, 1650–1655. doi:10.1126/science.1165594

Hervieux, N., Dumond, M., Sapala, A., Routier-Kierzkowska, A.-L., Kierzkowski, D., Roeder, A.H.K., Smith, R.S., Boudaoud, A., Hamant, O., 2016. A mechanical feedback restricts sepal growth and shape in arabidopsis. Curr. Biol. doi:10.1016/j.cub.2016.03.004

Hong, L., Dumond, M., Tsugawa, S., Sapala, A., Routier-Kierzkowska, A.-L., Zhou, Y., Chen, C., Kiss, A., Zhu, M., Hamant, O., Smith, R.S., Komatsuzaki, T., Li, C.-B., Boudaoud, A., Roeder, A.H.K., 2016. Variable Cell Growth Yields Reproducible OrganDevelopment through Spatiotemporal Averaging. Dev. Cell 38, 15–32. doi:10.1016/j.devcel.2016.06.016

Hong, L., Dumond, M., Zhu, M., Tsugawa, S., Li, C.-B., Boudaoud, A., Hamant, O., Roeder, A.H.K., 2018. Heterogeneity and robustness in plant morphogenesis: from cells to organs. Annu. Rev. Plant Biol. 69, 469–495. doi:10.1146/annurev-arplant-042817-040517

Jackson, M.D.B., Duran-Nebreda, S., Kierzkowski, D., Strauss, S., Xu, H., Landrein, B., Hamant, O., Smith, R.S., Johnston, I.G., Bassel, G.W., 2019. Global Topological Order Emerges through Local Mechanical Control of Cell Divisions in the Arabidopsis Shoot Apical Meristem. Cell Syst. 8, 53–65.e3. doi:10.1016/j.cels.2018.12.009

Jenik, P.D., Irish, V.F., 2000. Regulation of cell proliferation patterns by homeotic genes during *Arabidopsis* floral development. Development 127, 1267–1276.

Jürgens, G., Mayer, U., 1994. Arabidopsis, in: Bard, J. (Ed.), EMBRYOS, Colour Atlas of Developing Embryos. Wolfe Publishing, London, pp. 7–21.

Kierzkowski, D., Nakayama, N., Routier-Kierzkowska, A.-L., Weber, A., Bayer, E., Schorderet, M., Reinhardt, D., Kuhlemeier, C., Smith, R.S., 2012. Elastic domains regulate growth and organogenesis in the plant shoot apical meristem. Science 335, 1096–1099. doi:10.1126/science.1213100

Kierzkowski, D., Routier-Kierzkowska, A.-L., 2019. Cellular basis of growth in plants: geometry matters. Curr. Opin. Plant Biol. 47, 56–63. doi:10.1016/j.pbi.2018.09.008

Kierzkowski, D., Runions, A., Vuolo, F., Strauss, S., Lymbouridou, R., Routier-Kierzkowska, A.-L., Wilson-Sánchez, D., Jenke, H., Galinha, C., Mosca, G., Zhang, Z., Canales, C., Dello Ioio, R., Huijser, P., Smith, R.S., Tsiantis, M., 2019. A Growth-Based Framework for Leaf Shape Development and Diversity. Cell 177, 1405–1418.e17. doi:10.1016/j.cell.2019.05.011

Koncz, C., Schell, J., 1986. The promoter of TL-DNA gene 5 controls the tissue-specific expression of chimaeric genes carried by a novel Agrobacterium binary vector. Mol. Gen. Genet. 204, 383–396.

Kurihara, D., Mizuta, Y., Sato, Y., Higashiyama, T., 2015. ClearSee: a rapid optical clearing reagent for whole-plant fluorescence imaging. Development 142, 4168–4179. doi:10.1242/dev.127613

Landrein, B., Kiss, A., Sassi, M., Chauvet, A., Das, P., Cortizo, M., Laufs, P., Takeda, S., Aida, M., Traas, J., Vernoux, T., Boudaoud, A., Hamant, O., 2015. Mechanical stress contributes to the expression of the STM homeobox gene in Arabidopsis shoot meristems. elife 4, e07811. doi:10.7554/eLife.07811

Lee, K.J.I., Bushell, C., Koide, Y., Fozard, J.A., Piao, C., Yu, M., Newman, J., Whitewoods, C., Avondo, J., Kennaway, R., Marée, A.F.M., Cui, M., Coen, E., 2019. Shaping of a three-dimensional carnivorous trap through modulation of a planar growth mechanism. PLoS Biol. 17, e3000427. doi:10.1371/journal.pbio.3000427

Lein, E.S., Hawrylycz, M.J., Ao, N., Ayres, M., Bensinger, A., Bernard, A., Boe, A.F., Boguski, M.S., Brockway, K.S., Byrnes, E.J., Chen, Lin, Chen, Li, Chen, T.-M., Chin, M.C., Chong, J., Crook, B.E., Czaplinska, A., Dang, C.N., Datta, S., Dee, N.R., Jones, A.R., 2007. Genome-wide atlas of gene expression in the adult mouse brain. Nature 445, 168–176. doi:10.1038/nature05453

Liang, H., Mahadevan, L., 2011. Growth, geometry, and mechanics of a blooming lily. Proc Natl Acad Sci USA 108, 5516–5521. doi:10.1073/pnas.1007808108

Lieber, D., Lora, J., Schrempp, S., Lenhard, M., Laux, T., 2011. Arabidopsis WIH1 and WIH2 genes act in the transition from somatic to reproductive cell fate. Curr. Biol. 21, 1009–1017. doi:10.1016/j.cub.2011.05.015

Long, F., Peng, H., Liu, X., Kim, S.K., Myers, E., 2009. A 3D digital atlas of C. elegans and its application to single-cell analyses. Nat. Methods 6, 667–672. doi:10.1038/nmeth.1366

Lora, J., Herrero, M., Tucker, M.R., Hormaza, J.I., 2017. The transition from somatic to germline identity shows conserved and specialized features during angiosperm evolution. New Phytol. 216, 495–509. doi:10.1111/nph.14330

Louveaux, M., Julien, J.-D., Mirabet, V., Boudaoud, A., Hamant, O., 2016. Cell division plane orientation based on tensile stress in *Arabidopsis thaliana*. Proc Natl Acad Sci USA 113, E4294–303. doi:10.1073/pnas.1600677113

Mansfield, S.G., Briarty, L.G., Erni, S., 1991. Early embryogenesis in *Arabidopsis thaliana*. I. The mature embryo sac. Can J Bot 69, 447–460.

Mayer, K.F., Schoof, H., Haecker, A., Lenhard, M., Jürgens, G., Laux, T., 1998. Role of WUSCHEL in regulating stem cell fate in the Arabidopsis shoot meristem. Cell 95, 805–815. doi:10.1016/s0092-8674(00)81703-1

Meister, R.J., Kotow, L.M., Gasser, C.S., 2002. SUPERMAN attenuates positive INNER NO OUTER autoregulation to maintain polar development of Arabidopsis ovule outer integuments. Development 129, 4281–4289.

Meister, R.J., Williams, L.A., Monfared, M.M., Gallagher, T.L., Kraft, E.A., Nelson, C.G., Gasser, C.S., 2004. Definition and interactions of a positive regulatory element of the Arabidopsis INNER NO OUTER promoter. Plant J. 37, 426–438.

Montenegro-Johnson, T.D., Stamm, P., Strauss, S., Topham, A.T., Tsagris, M., Wood, A.T.A., Smith, R.S., Bassel, G.W., 2015. Digital Single-Cell Analysis of Plant Organ Development Using 3DCellAtlas. Plant Cell 27, 1018–1033. doi:10.1105/tpc.15.00175

Montenegro-Johnson, T., Strauss, S., Jackson, M.D.B., Walker, L., Smith, R.S., Bassel, G.W., 2019. 3DCellAtlas Meristem: a tool for the global cellular annotation of shoot apical meristems. Plant Methods 15, 33. doi:10.1186/s13007-019-0413-0

Musielak, T.J., Schenkel, L., Kolb, M., Henschen, A., Bayer, M., 2015. A simple and versatile cell wall staining protocol to study plant reproduction. Plant Reprod. 28, 161–169. doi:10.1007/s00497-015-0267-1

Nakajima, K., 2018. Be my baby: patterning toward plant germ cells. Curr. Opin. Plant Biol. 41, 110–115. doi:10.1016/j.pbi.2017.11.002

Oliphant, T.E., 2007. Python for Scientific Computing. Comput. Sci. Eng. 9, 10–20. doi:10.1109/MCSE.2007.58

Pasternak, T., Haser, T., Falk, T., Ronneberger, O., Palme, K., Otten, L., 2017. A 3D digital atlas of the Nicotiana tabacum root tip and its use to investigate changes in the root apical meristem induced by the Agrobacterium 6b oncogene. Plant J. 92, 31–42. doi:10.1111/tpj.13631

Rebocho, A.B., Southam, P., Kennaway, J.R., Bangham, J.A., Coen, E., 2017. Generation of shape complexity through tissue conflict resolution. elife 6. doi:10.7554/eLife.20156

Rein, K., Zöckler, M., Mader, M.T., Grübel, C., Heisenberg, M., 2002. The Drosophila standard brain. Curr. Biol. 12, 227–231. doi:10.1016/s0960-9822(02)00656-5

Robinson-Beers, K., Pruitt, R.E., Gasser, C.S., 1992. Ovule development in wild-Type Arabidopsis and two female-sterile mutants. Plant Cell 4, 1237–1249. doi:10.1105/tpc.4.10.1237

Ronneberger, O., Fischer, P., Brox, T., 2015. U-Net: Convolutional Networks for Biomedical Image Segmentation, in: Navab, N., Hornegger, J., Wells, W.M., Frangi, A.F. (Eds.), Medical Image Computing and Computer-Assisted Intervention (MICCAI), Lecture Notes in Computer Science. Springer International Publishing, Cham, pp. 234–241. doi:10.1007/978-3-319-24574-4_28

RStudio Team, 2019. RStudio: Integrated Development for R. RStudio, Inc., Boston, MA, USA.

R Core Team, 2019. R: A language and environment for statistical computing. R Foundation for Statistical Computing, Vienna.

Sampathkumar, A., Krupinski, P., Wightman, R., Milani, P., Berquand, A., Boudaoud, A., Hamant, O., Jönsson, H., Meyerowitz, E.M., 2014. Subcellular and supracellular mechanical stress prescribes cytoskeleton behavior in Arabidopsis cotyledon pavement cells. elife 3, e01967. doi:10.7554/eLife.01967

Sankar, M., Nieminen, K., Ragni, L., Xenarios, I., Hardtke, C.S., 2014. Automated quantitative histology reveals vascular morphodynamics during Arabidopsis hypocotyl secondary growth. elife 3, e01567. doi:10.7554/eLife.01567

Sapala, A., Runions, A., Routier-Kierzkowska, A.-L., Das Gupta, M., Hong, L., Hofhuis, H., Verger, S., Mosca, G., Li, C.-B., Hay, A., Hamant, O., Roeder, A.H., Tsiantis, M., Prusinkiewicz, P., Smith, R.S., 2018. Why plants make puzzle cells, and how their shape emerges. elife 7. doi:10.7554/eLife.32794

Sapala, A., Runions, A., Smith, R.S., 2019. Mechanics, geometry and genetics of epidermal cell shape regulation: different pieces of the same puzzle. Curr. Opin. Plant Biol. 47, 1–8. doi:10.1016/j.pbi.2018.07.017

Sarkans, U., Gostev, M., Athar, A., Behrangi, E., Melnichuk, O., Ali, A., Minguet, J., Rada, J.C., Snow, C., Tikhonov, A., Brazma, A., McEntyre, J., 2018. The BioStudies database-one stop shop for all data supporting a life sciences study. Nucleic Acids Res. 46, D1266–D1270. doi:10.1093/nar/gkx965

Sassi, M., Ali, O., Boudon, F., Cloarec, G., Abad, U., Cellier, C., Chen, X., Gilles, B., Milani, P., Friml, J., Vernoux, T., Godin, C., Hamant, O., Traas, J., 2014. An auxin-mediated shift toward growth isotropy promotes organ formation at the shoot meristem in *Arabidopsis*. Curr. Biol. 24, 2335–2342. doi:10.1016/j.cub.2014.08.036

Satina, S., Blakeslee, A.F., Avery, A.G., 1940. Demonstration of the three germ layers in the shoot apex of Datura by means of induced polyploidy in periclinal chimeras. Am J Bot 27, 895–905.

Schmidt, A., Schmid, M.W., Grossniklaus, U., 2015. Plant germline formation: common concepts and developmental flexibility in sexual and asexual reproduction. Development 142, 229–241. doi:10.1242/dev.102103

Schmidt, T., Pasternak, T., Liu, K., Blein, T., Aubry-Hivet, D., Dovzhenko, A., Duerr, J., Teale, W., Ditengou, F.A., Burkhardt, H., Ronneberger, O., Palme, K., 2014. The iRoCS Toolbox--3D analysis of the plant root apical meristem at cellular resolution. Plant J. 77, 806–814. doi:10.1111/tpj.12429

Schneitz, K., Hülskamp, M., Kopczak, S.D., Pruitt, R.E., 1997. Dissection of sexual organ ontogenesis: a genetic analysis of ovule development in *Arabidopsis thaliana*. Development 124, 1367–1376.

Schneitz, K., Hülskamp, M., Pruitt, R.E., 1995. Wild-type ovule development in *Arabidopsis thaliana*: a light microscope study of cleared whole-mount tissue. Plant J. 7, 731–749. doi:10.1046/j.1365-313X.1995.07050731.x

Sieber, P., Gheyselinck, J., Gross-Hardt, R., Laux, T., Grossniklaus, U., Schneitz, K., 2004. Pattern formation during early ovule development in *Arabidopsis thaliana*. Dev. Biol. 273, 321–334. doi:10.1016/j.ydbio.2004.05.037

Sladitschek, H.L., Fiuza, U.-M., Pavlinic, D., Benes, V., Hufnagel, L., Neveu, P.A., 2020. MorphoSeq: full single-cell transcriptome dynamics up to gastrulation in a chordate. Cell 181, 922–935.e21. doi:10.1016/j.cell.2020.03.055

Speth, M., Andres, B., Reinelt, G., Schnörr, C., 2011. Globally optimal image partitioning by multicuts. Presented at the International Workshop on Energy Minimization Methods in Computer Vision and Pattern Recognition, Springer, pp. 31–44.

Strauss, S., Sapala, A., Kierzkowski, D., Smith, R.S., 2019. Quantifying Plant Growth and Cell Proliferation with MorphoGraphX. Methods Mol. Biol. 1992, 269–290. doi:10.1007/978-1-4939-9469-4_18

Tofanelli, R., Vijayan, A., Scholz, S., Schneitz, K., 2019. Protocol for rapid clearing and staining of fixed Arabidopsis ovules for improved imaging by confocal laser scanning microscopy. Plant Methods 15, 120. doi:10.1186/s13007-019-0505-x

Truernit, E., Haseloff, J., 2008. Arabidopsis thaliana outer ovule integument morphogenesis: ectopic expression of KNAT1 reveals a compensation mechanism. BMC Plant Biol. 8, 35. doi:10.1186/1471-2229-8-35

Tucker, M.R., Okada, T., Hu, Y., Scholefield, A., Taylor, J.M., Koltunow, A.M.G., 2012. Somatic small RNA pathways promote the mitotic events of megagametogenesis during female reproductive development in Arabidopsis. Development 139, 1399–1404. doi:10.1242/dev.075390

Uyttewaal, M., Burian, A., Alim, K., Landrein, B., Borowska-Wykręt, D., Dedieu, A., Peaucelle, A., Ludynia, M., Traas, J., Boudaoud, A., Kwiatkowska, D., Hamant, O., 2012. Mechanical stress acts via katanin to amplify differences in growth rate between adjacent cells in Arabidopsis. Cell 149, 439–451. doi:10.1016/j.cell.2012.02.048

Valuchova, S., Mikulkova, P., Pecinkova, J., Klimova, J., Krumnikl, M., Bainar, P., Heckmann, S., Tomancak, P., Riha, K., 2020. Imaging plant germline differentiation within Arabidopsis flowers by light sheet microscopy. elife 9. doi:10.7554/eLife.52546

Van Hooijdonk, C.A., Glade, C.P., Van Erp, P.E., 1994. TO-PRO-3 iodide: a novel HeNe laser-excitable DNA stain as an alternative for propidium iodide in multiparameter flow cytometry. Cytometry 17, 185–189. doi:10.1002/cyto.990170212

Villanueva, J.M., Broadhvest, J., Hauser, B.A., Meister, R.J., Schneitz, K., Gasser, C.S., 1999. INNER NO OUTER regulates abaxial-adaxial patterning in Arabidopsis ovules. Genes Dev. 13, 3160–3169.

Wang, Z.-P., Xing, H.-L., Dong, L., Zhang, H.-Y., Han, C.-Y., Wang, X.-C., Chen, Q.-J., 2015. Egg cell-specific promoter-controlled CRISPR/Cas9 efficiently generates homozygous mutants for multiple target genes in Arabidopsis in a single generation. Genome Biol. 16, 144. doi:10.1186/s13059-015-0715-0

Wolny, A., Cerrone, L., Vijayan, A., Tofanelli, R., Barro, A.V., Louveaux, M., Wenzl, C., Strauss, S., Wilson-Sánchez, D., Lymbouridou, R., Steigleder, S., Pape, C., Bailoni, A., Duran-Nebreda, S., Bassel, G., Lohmann, J.U., Tsiantis, M., Hamprecht, F., Schneitz, K., Maizel, A., Kreshuk, A., 2020. Accurate and versatile 3D segmentation of plant tissues at cellular resolution. elife 9. doi:10.7554/eLife.57613

Xie, K., Zhang, J., Yang, Y., 2014. Genome-wide prediction of highly specific guide RNA spacers for CRISPR-Cas9-mediated genome editing in model plants and major crops. Mol. Plant 7, 923–926. doi:10.1093/mp/ssu009

Yoshida, S., Barbier de Reuille, P., Lane, B., Bassel, G.W., Prusinkiewicz, P., Smith, R.S., Weijers, D., 2014. Genetic control of plant development by overriding a geometric division rule. Dev. Cell 29, 75–87. doi:10.1016/j.devcel.2014.02.002

Zhao, X., Bramsiepe, J., Van Durme, M., Komaki, S., Prusicki, M.A., Maruyama, D., Forner, J., Medzihradszky, A., Wijnker, E., Harashima, H., Lu, Y., Schmidt, A., Guthörl, D., Logroño, R.S., Guan, Y., Pochon, G., Grossniklaus, U., Laux, T., Higashiyama, T., Lohmann, J.U., Schnittger, A., 2017. RETINOBLASTOMA RELATED1 mediates germline entry in Arabidopsis. Science 356. doi:10.1126/science.aaf6532

